# Diploid hepatocytes drive physiological liver renewal in adult humans

**DOI:** 10.1101/2020.08.07.230086

**Authors:** Paula Heinke, Fabian Rost, Julian Rode, Thilo Welsch, Kanar Alkass, Joshua Feddema, Mehran Salehpour, Göran Possnert, Henrik Druid, Lutz Brusch, Olaf Bergmann

## Abstract

Physiological liver cell replacement is central to maintaining the organ’s high metabolic activity, although its characteristics are difficult to study in humans. Using retrospective ^14^C birth dating of cells, we report that human hepatocytes show continuous and lifelong turnover, maintaining the liver a young organ (average age < 3 years). Hepatocyte renewal is highly dependent on the ploidy level. Diploid hepatocytes show an seven-fold higher annual exchange rate than polyploid hepatocytes. These observations support the view that physiological liver cell renewal in humans is mainly dependent on diploid hepatocytes, whereas polyploid cells are compromised in their ability to divide. Moreover, cellular transitions between these two subpopulations are limited, with minimal contribution to the respective other ploidy class under homeostatic conditions. With these findings, we present a new integrated model of homeostatic liver cell generation in humans that provides fundamental insights into liver cell turnover dynamics.

## Introduction

The liver has a remarkable potential to generate new functional tissue in response to injury, which relies upon the proliferative capacity of liver cells. However, in humans, it remains unknown whether parenchymal hepatocytes are constantly exchanged throughout one’s lifetime or whether they are physiologically long-lived cells, similar to cardiomyocytes and neurons (Bergmann et al., 2015; Huttner et al., 2014), that maintain structural and functional integrity over the span of several decades.

During both homeostasis and disease, the replacement of liver parenchyma relies on hepatocytes (Malato et al., 2011; Schaub et al., 2014; Wang et al., 2017; Yanger et al., 2014). The contribution of other cells seems to be only relevant in cases of severe chronic damage when the normal hepatocyte response is exhausted (Deng et al., 2018; Lu et al., 2015; Raven et al., 2017).

Reports in recent decades, which have relied mainly on cell cycle markers and nucleotide analogs, have estimated the age of hepatocytes to be between 200 and 400 days in rodents (Macdonald, 1961; Magami et al., 2002). On the other hand, a recent pulse-chase study applying advanced isotope labeling supported the view that within the life span of a mouse, parenchymal liver cells are only rarely exchanged and most are as old as neurons (Arrojo e Drigo et al., 2019). Estimations of hepatocyte turnover in humans are scarce due to strict methodological limitations. Therefore, it is not known whether the reported hepatocyte renewal paradigm in mice can translate to a human life span of more than 80 years.

Furthermore, it is unclear how pronounced hepatocyte heterogeneity affects cellular turnover. Recent animal studies have provided evidence of distinct subpopulations with special proliferative capacities. Reports have shown that hepatocytes from both the periportal region (Font-Burgada et al., 2015; Pu et al., 2016) and the pericentral region (Wang et al., 2015), as well as randomly distributed hepatocytes (Lin et al., 2018; Planas-Paz et al., 2016), are responsible for maintaining the hepatocyte pool. However, new studies have challenged these findings, and the existence of specific proliferative subpopulations characterized by their zonation has been questioned (Chen et al., 2020; Sun et al., 2020).

In contrast to most other cells in the human body, a fraction of hepatocytes acquire a polyploid state during normal aging (Kudryavtsev et al., 1993). Polyploidy has been attributed to cell cycle arrest and restricted proliferation in the liver and other tissues (Derks and Bergmann, 2020; Donne et al., 2020; Ganem et al., 2014; Wilkinson et al., 2019). This property has been questioned with regard to hepatocytes based on the results of several mouse studies that have described an equivalent proliferative capacity irrespective of their ploidy level (Duncan et al., 2010; Zhang et al., 2018). Importantly, the evolution and degree of polyploidization show significant interspecies differences. Sixty to seventy percent of human hepatocytes remain diploid over a lifetime (Duncan et al., 2012; Kudryavtsev et al., 1993), while up to ninety percent increase their ploidy level in mice (Duncan et al., 2010; Wang et al., 2014). However, the reasons and biological consequences of these variations are poorly understood.

To date, a comprehensive exploration characterizing the dynamics of human hepatocyte turnover, including their age distribution and the consequences for hepatocyte functionality in the aging liver, is lacking. It is important to establish these characteristics of cell renewal in the adult liver, particularly to gain a better understanding of age-related diseases and the development of liver cancer. Here, we explore the dynamics of human liver cell turnover and the age distribution of hepatocytes through retrospective ^14^C birth dating and mathematical modeling. We provide a comprehensive model describing liver cell turnover in humans.

## Results

### Liver nucleus isolation for retrospective ^14^C birth dating

Nuclei were isolated from human liver tissue using density gradient centrifugation and incubated with an antibody against the nuclear transcription factor hepatocyte nuclear factor 4 alpha (HNF4α). HNF4α is specifically expressed in hepatocyte nuclei (Si-Tayeb et al., 2010) and therefore enables their identification and separation from nonhepatocyte nuclei isolated from fresh or frozen liver tissue (Fig. 1A, B). Using FACS, we isolated 73.6% ± 10.5% (mean ± SD) hepatocytes from purified nuclear fractions with no age-related changes (Fig. 1B and S1B) in accordance with the known cellular composition of the human liver (Donne et al., 2020). Purities of the sorted HNF4α-positive hepatocyte (98.2% ± 0.9%) and HNF4α-negative nonhepatocyte nuclei populations (97.3% ± 1.5%) were verified by FACS reanalyses (Fig. S1A).

**Figure 1:**
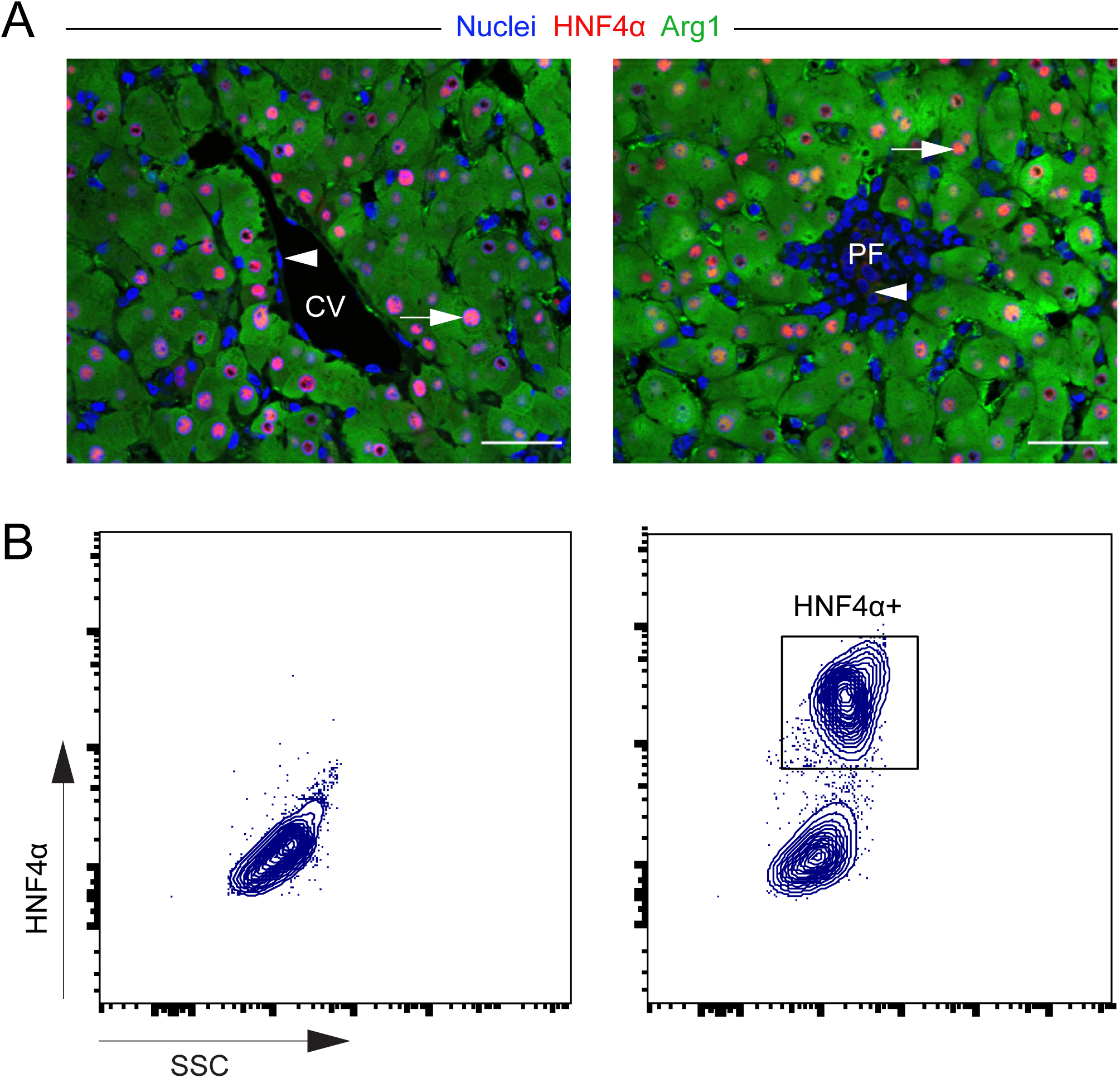
Identification and separation of hepatocyte nuclei in human liver tissue. (A) HNF4α is specifically expressed in all hepatocyte nuclei (arrows) but not in nonhepatocytes (arrowheads) (CV = central vein, PF = portal field), documented by colabeling with an established cytoplasmic marker for hepatocytes (Arg1), scale bar 50 µm. (B) HNF4α immunolabeling (right) can be used to isolate hepatocyte nuclei from nonhepatocyte nuclei using FACS compared with a negative control (left).

### Retrospective ^14^C birth dating of human liver cells

Genomic DNA was extracted from 34 subjects, aged 20-84 (Table S1), from FACS-isolated hepatocyte nuclei (n = 31), nonhepatocyte nuclei (n = 11) and unsorted liver nuclei populations (n = 23). ^14^C concentrations were determined using accelerator mass spectrometry (Bergmann et al., 2015; Spalding et al., 2013) (Table S2). We corrected ^14^C concentrations for impurities introduced during nucleus sorting, as described (see methods). By comparing the measured ^14^C values of the samples to historic ^14^C atmospheric levels, we estimated the average genomic age of hepatic cells (Fig. 2A and Fig S2A). The results indicate a continuous and substantial turnover over the entire lifetime with an average genomic ^14^C age of 6.1 ± 3.9 years, independent of subject age (Fig. S2B). This finding was confirmed by genomic ^14^C concentrations of sorted hepatocyte nuclei (Fig. 2B, C) and nonhepatocyte nuclei (Fig. S2C, D) with average ^14^C ages of 4.4 ± 4.3 years and 8.8 ± 7.0 years, respectively. This result suggests that the human liver is a highly regenerative organ, with most cells being continuously replaced over the human life span.

**Figure 2:**
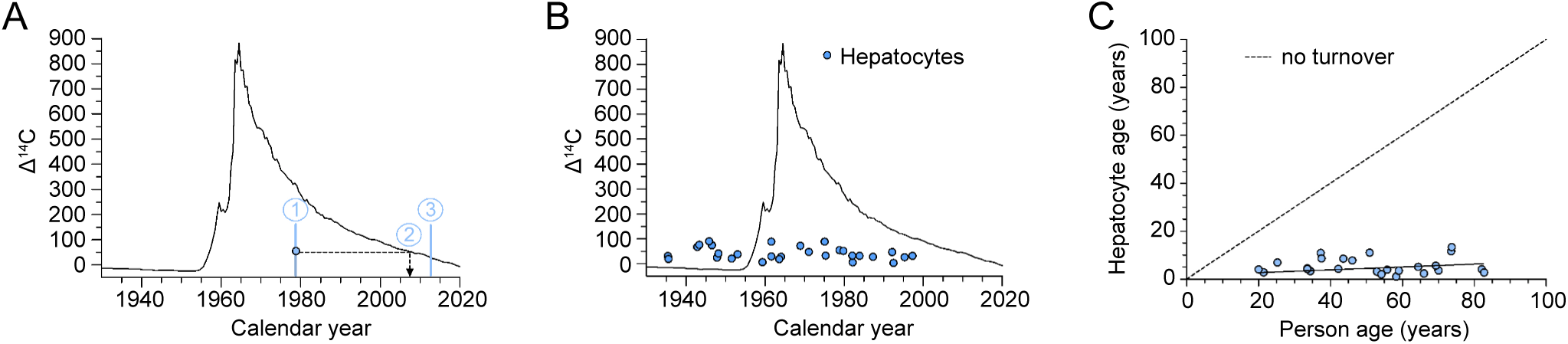
^14^C retrospective birth dating of human hepatocytes. (A) Schematic representation of ^14^C data analysis. Measured ^14^C data (blue dots) are depicted at the persons’ birth date (1). The curve of the equivalent level of the atmospheric ^14^C (black line) would represent the time point of ^14^C formation of the analyzed cell population (2, dashed line) if all cells were born at the same time. The difference between the time of sample acquisition (3) and the time point of ^14^C formation (2) provides an estimate of the genomic carbon age. (B) Genomic ^14^C concentrations of hepatocyte nuclei do not correlate with atmospheric ^14^C concentration at the time of birth. (C) The estimated carbon age of hepatocyte nuclei is independent of the persons’ age, indicating continuous lifelong cellular turnover (n = 29, R = 0.26, p = 0.18).

### Dynamics of hepatocyte renewal in humans

To characterize the generation of human hepatocytes from our measured genomic ^14^C data, we used a mathematical modeling strategy (Bergmann et al., 2015; Huttner et al., 2014; Spalding et al., 2013) (see supplementary methods). We derived a population balance equation for ^14^C concentration structured cell populations, which we used to predict the mean ^14^C concentration dynamics from cell turnover rates. We constrained the turnover rates in such a way that the total DNA content of the liver remained constant. Using an additive Gaussian error model, we then estimated the cell turnover rates using Bayesian inference. Different population structures and turnover dynamics translate to different model variants, which we termed scenarios. We selected the best-fitting scenario, i.e., the scenario with the highest predictive power, using the leave-one-out (LOO) cross-validation method (Vehtari et al., 2017).

With this approach, we first developed a scenario to analyze the rate at which hepatocytes renew in general, assuming that all cells are replaced at the same frequency. This one-population scenario (POP1) predicts that hepatocytes are exchanged at a rate of 20% per year, with no evidence of an age-dependent change in turnover (Fig. S4A, B and supplementary methods).

Hepatocytes that would become post-mitotic soon after a person’s birth had a ^14^C concentration corresponding to the atmospheric level at that time point, thus allowing the existence of a nondividing quiescent subpopulation to be tested (scenario POP1q, Fig. S4A and supplementary methods). The POP1q scenario did not fit the data better than the one-population model described above and predicted that the vast majority of the hepatocyte population (> 96%) is renewed with an annual turnover rate of 25%. Thus, there is no evidence of the existence of a considerable subpopulation of quiescent, long-lived hepatocytes.

With our model, we further considered the possibility of two independent hepatocyte subpopulations with different turnover dynamics (scenario POP2, Fig. S4A and supplementary methods). We found that if these two populations existed, they would have a ratio of 37% vs. 63%, and the hepatocytes would renew at very similar rates as in the above described one-population model (16% and 23% per year, respectively). These findings do not support the presence of independent hepatocyte subpopulations with distinctly different turnover dynamics.

### Hepatocyte ploidy increases over the lifetime in humans

Rodents develop a considerable amount of hepatocyte binucleation and polyploidy starting around the time of weaning (Guidotti et al., 2003; Margall-Ducos et al., 2007). In contrast, polyploidization events in humans occur much later in life, beginning at approximately 50 years of age and gradually increasing the overall ploidy in the aged liver (Kudryavtsev et al., 1993). We applied quantitative imaging and flow cytometry to describe the course of hepatocyte binucleation and polyploidization in human liver tissue samples (Fig. 3A, C and Fig. S3A, B). We observed substantial changes in the cellular and nuclear ploidy composition over the lifetime (Fig. 3B, D), with the number of diploid hepatocytes decreasing with age, in line with previous studies (Kudryavtsev et al., 1993). While more than 80% of hepatocytes are still diploid in young adults (20-45 years), this fraction drops to less than 70% in old subjects (> 65 years) (Fig. 3B) (Kudryavtsev et al., 1993). Accordingly, the percentage of polyploid hepatocytes, particularly binucleated diploid and mononucleated tetraploid, increases steadily over the lifetime, reaching up to 35% of all hepatocytes at advanced ages (Fig. S3C). Higher ploidy classes represent less than 10% of all hepatocytes in adulthood, with less than 5% in young and middle-aged (45-65 years) individuals.

**Figure 3:**
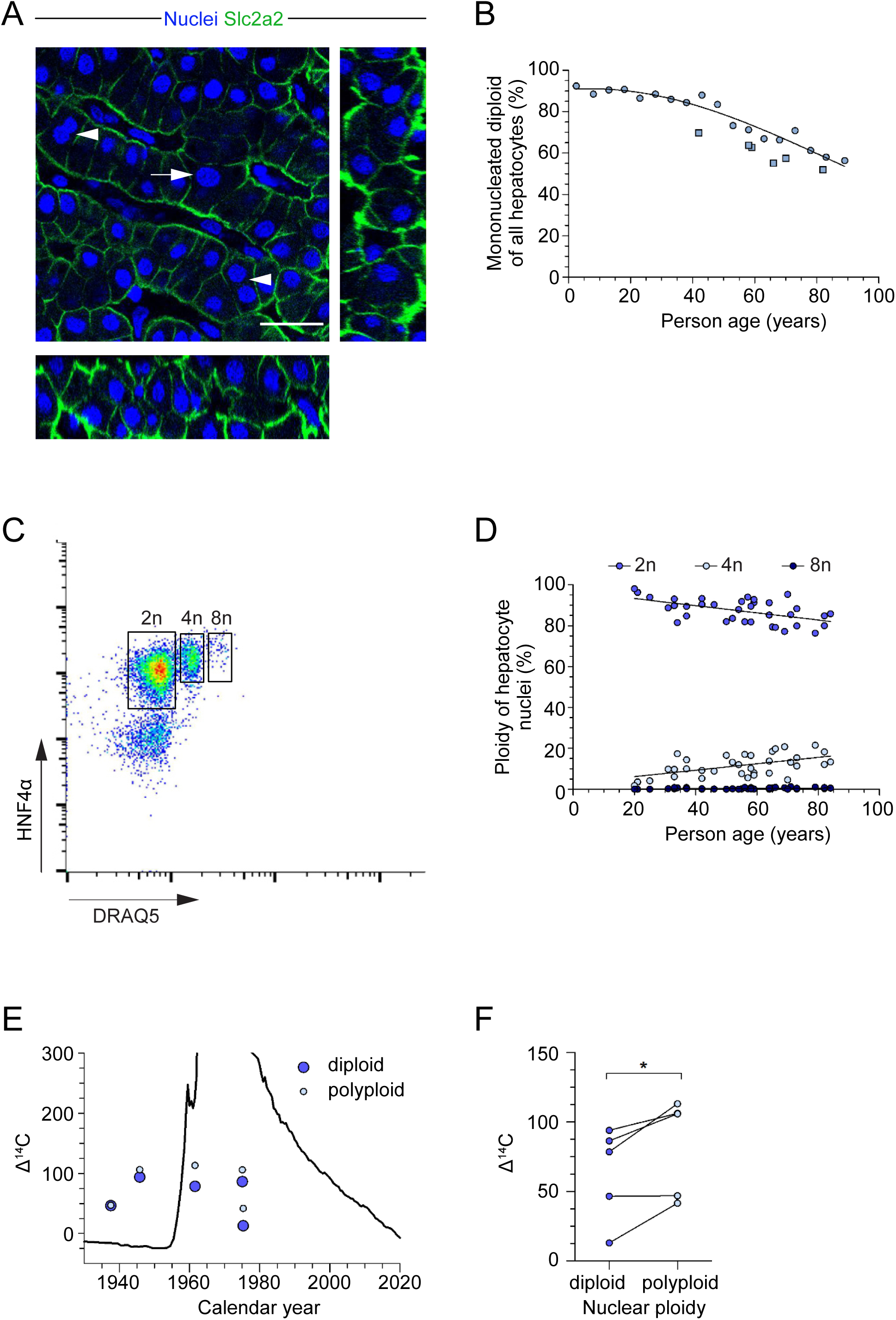
The human liver is composed of polyploid hepatocytes with different genomic ^14^C ages. (A) Human hepatocytes exhibit cellular (arrowheads) and nuclear (arrow) ploidy, scale bar 25 µm. (B) Quantitative image analysis of hepatocytes confirms an age-dependent decrease of mononucleated, diploid hepatocytes (squares), as described previously by Kudryavtsev et al., 1993 (circles) (Gaussian regression, n = 161, R = 0.81). (C) Separation of diploid and polyploid hepatocyte nuclei with DNA stain (DRAQ5) using FACS. (D) Diploid hepatocyte nuclei decrease with age (linear regression, n = 35, R = 0.54, p = 0.0008), while the fraction of tetraploid nuclei increases (linear regression, n = 35, R = 0.53, p = 0.0010). Octaploid hepatocyte nuclei represent only a small fraction of hepatocyte nuclei in the adult liver (linear regression, n = 35, R = 0.36, p = 0.04). (E and F) Polyploid hepatocyte nuclei show higher genomic ^14^C levels than diploid nuclei (paired t-test, * p = 0.03).

### Ploidy is associated with differences in the age and renewal rates of human hepatocytes

We analyzed the genomic ^14^C concentrations of hepatocyte nuclei based on different nuclear ploidy levels, extending the flow cytometry sorting strategy described above (Fig. 3E). Reanalyses confirmed the purity of the sorted diploid and polyploid populations (Fig. S3D). Analysis of five different cases revealed that polyploid hepatocyte nuclei show significantly higher ^14^C concentrations (ΔΔ^14^C = 19.1 ± 12.1) than diploid hepatocyte nuclei from the same individual (Fig. 3F). The ^14^C concentrations of polyploid hepatocyte nuclei correspond to time points before the ^14^C formation date of diploid hepatocyte nuclei, suggesting that polyploid hepatocytes are older than diploid hepatocytes and might have different turnover characteristics (Fig. 2A and Fig. S3E).

Consequently, we developed a more refined mathematical scenario (scenario POP2p), describing a diploid and a polyploid hepatocyte population with distinct rates of cell division and cell death (Fig. 4A, S4A and supplementary methods). This scenario also accounts for ploidy exchanges between the diploid and polyploid populations. We constrained the turnover rates such that the scenario correctly predicted the changes in the composition of ploidy populations during adulthood (Fig. 3B, D, 4A and supplementary methods). Because most polyploid hepatocytes are tetraploid (Fig. S3C), we neglected higher ploidy levels in this scenario, and assumed that binucleated diploid (2×2n) and mononucleated tetraploid cells (4n) behave identically.

**Figure 4:**
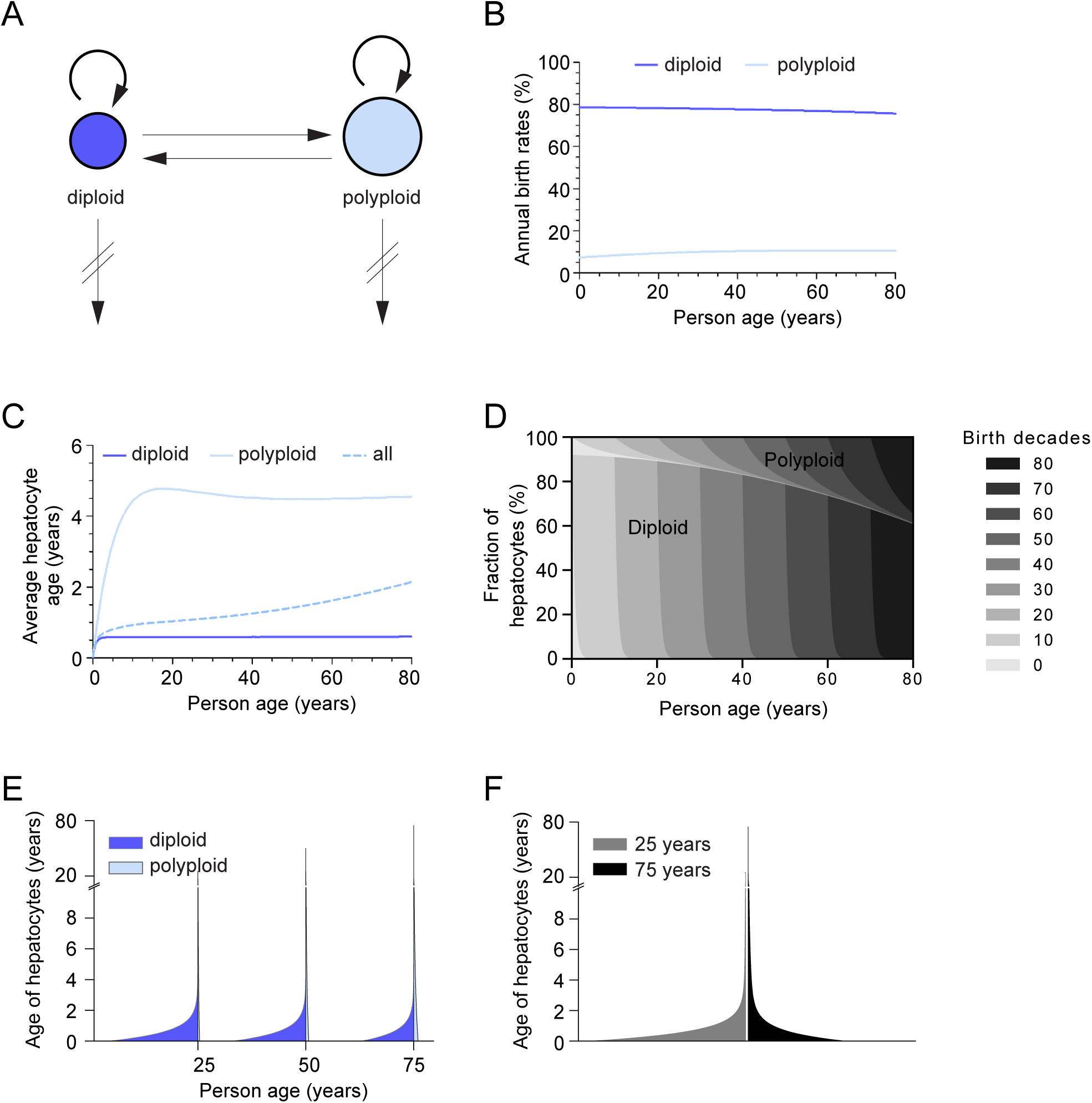
Ploidy is associated with the renewal dynamics and age distribution of human hepatocytes. (A) The mathematical scenario POP2p with the best fit to our measured genomic ^14^C levels describes cellular birth (circular arrows) and death rates (downwards arrows) of diploid and polyploid (including tetraploid and binuclear) hepatocytes and an exchange between the ploidy classes (horizontal arrows). (B) Estimated annual birth rates of diploid and polyploid hepatocytes remain nearly constant over a person’s lifetime. (C) The average cellular age of polyploid hepatocytes is more than seven-fold higher than that of diploid hepatocytes. Because polyploidization advances with age, the overall hepatocyte age increases. (D) The age distribution of human hepatocytes is illustrated for a person’s given age. The decade in which a cell was born is shown in different shades of gray. Polyploid hepatocytes remain in the liver much longer than diploid hepatocytes. (E) Age distribution of diploid and polyploid hepatocytes at 25, 50 and 75 years of age. The average cellular age as well as the fraction of polyploid cells increases within the lifetime. (F) Comparative age distribution of all hepatocytes at 25 and 75 years of age illustrates the accumulation of older cells over the lifetime.

This scenario fits the measured ^14^C data best among all scenarios described above because it accounted for the differences in the genomic ^14^C concentrations in diploid and polyploid hepatocytes (Fig. S4C and supplementary methods). In the POP2p scenario, the renewal dynamics of diploid and polyploid hepatocytes showed no considerable age-related changes, in line with the above tested scenarios (Fig. 4B). However, diploid and polyploid hepatocyte populations display very different turnover characteristics. In a middle-aged individual, the annual cell division rate for diploid hepatocytes is approximately 77%, whereas it is only 11% among polyploid hepatocytes (Fig. 4B), corresponding to average cell ages of 0.6 years for diploid hepatocytes and 4.5 years for polyploid hepatocytes (Fig. 4C). Accordingly, we estimate that approximately 175 million hepatocytes are born every day in a human middle-aged liver (see supplementary methods). While almost all diploid cells are exchanged within one year, up to 12% of polyploid cells reside more than one decade in the organ (Fig. 4D). As the fraction of polyploid hepatocytes increases over the lifetime, the age distribution of hepatocytes changes accordingly in aging individuals (Fig. 4D-F). In a 25-year-old individual, 89% of hepatocytes are less than two years old, whereas this fraction decreases to 75% in a 75-year-old subject. Although polyploid hepatocytes live substantially longer than diploid hepatocytes, most liver cells are short-lived, irrespective of age, and more than half of all hepatocytes have been born within the last year.

### The ploidy conveyer in human liver

Our best-fitting scenario (scenario POP2p) not only estimates the dynamics of hepatocyte generation for diploid and polyploid cells but also describes cellular transitions between ploidy classes, as previously suggested with the concept of a ploidy conveyer (Duncan et al., 2010) (Fig. 4A). According to our model, this bidirectional exchange between hepatocyte populations is necessary to explain the measured ^14^C age distributions. In a 25-year-old subject, approximately 17% of polyploid hepatocytes arise from diploid cells (Fig. 5A). Due to the continuous increase in ploidy level, this exchange rate drops to 4% in a 75-year-old individual. In contrast, only 1% of the diploid population originates from polyploid hepatocytes at a young age, and 4% of diploid hepatocytes are generated by the polyploid hepatocyte pool in old individuals (Fig. 5B). However, the vast majority of diploid and polyploid hepatocytes are maintained through renewal processes within their individual ploidy class (Fig. 5A, B).

**Figure 5:**
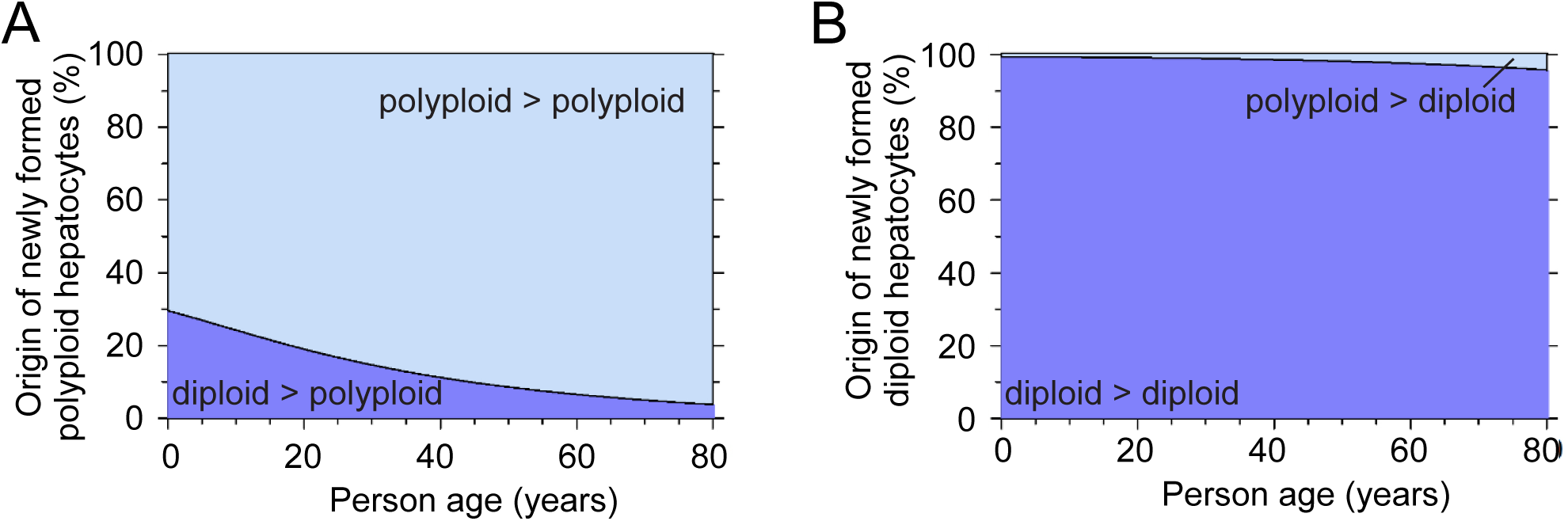
Ploidy conveyer in human hepatocytes. (A) There is a lifelong contribution of diploid hepatocytes to the polyploid cell pool. However, the majority of polyploid hepatocytes arise from cells of the same ploidy class by cell division. (B)Most diploid hepatocytes originate from the pool of diploid hepatocytes, and only a small fraction originates from the polyploid hepatocyte pool by ploidy reversal.

## Discussion

Our current understanding of human liver homeostasis is far from complete, and we have limited knowledge regarding to what degree the replacement of parenchymal cells is required to maintain the high metabolic activity of hepatocytes. Here, we used ^14^C retrospective birth dating to evaluate the birth rate and turnover dynamics of human liver cells. We found fundamental differences in the renewal between diploid and polyploid hepatocytes: diploid cells are exchanged approximately once a year, while higher ploidy classes reside up to one decade in the organ.

### The human liver remains a young organ

It is generally agreed that hepatocytes retain their ability to renew in adulthood under physiological conditions (Macdonald, 1961). A clear picture of the renewal behavior of parenchymal liver cells in humans has been lacking until now. The turnover rate of adult hepatocytes has been studied in a number of animal models (Arrojo e Drigo et al., 2019; Magami et al., 2002), but it remains unknown whether these results can be translated to humans. A recent study using ^15^N labeling showed that the majority of murine hepatocytes are not replaced within the observational period of 18 months (Arrojo e Drigo et al., 2019). The authors concluded that most hepatocytes are not exchanged and are as old as post-mitotic neurons. However, in humans, one and a half years represents a rather short episode in life, and much longer observation periods are necessary to conclude that most hepatocytes do not renew in a lifetime. Studies in humans, which are based on the expression of the cell cycle marker Ki-67, have revealed a very low labeling index in homeostasis (< 0.5%) (Vansaun et al., 2013) (Fig. S1C). However, such imaging approaches reflect cell cycle activity only at a single time point without knowing the fate of the cycling cell. Moreover, they do not provide information as to whether there are subpopulations of hepatocytes that show different cycling frequencies. Thus, these studies based on the detection of cell cycle markers are insufficient to establish reliable turnover rates in humans (Spalding et al., 2008).

In contrast, ^14^C birth dating provides a cumulative measure of cell renewal, which makes it possible to retrospectively study cellular turnover in human tissue samples (Spalding et al., 2005). ^14^C birth dating has been applied to many organs to characterize cellular ages and turnover dynamics (Bergmann et al., 2009; Spalding et al., 2013; Yeung et al., 2019). These studies have shown that ^14^C levels in genomic DNA accurately represent atmospheric ^14^C concentrations from the time point of cell generation. Exchange of carbon atoms in genomic DNA at later time points, such as methylation and DNA repair, does not affect the measured ^14^C levels (Spalding et al., 2005). Recently, independent laboratories using next-generation sequencing observed low amounts (< 5%) of aneuploidy and copy number variations in hepatocytes (Knouse et al., 2014; Sladky et al., 2020). These studies suggest that the genomic stability of hepatocytes is similar to that of neurons (Knouse et al., 2014) and therefore would not affect the results of ^14^C retrospective birth dating (Bergmann et al., 2012; Huttner et al., 2014).

We applied retrospective ^14^C birth dating to explore cell turnover in the human liver. By analyzing unsorted liver nuclei, we report an average liver cell age of less than three years using the simplified one-population scenario (POP1, Fig. S4A). This estimation considers a semiconservative replication mechanism and therefore predicts cellular ages that are lower than the measured genomic ^14^C age (Fig. 2A). This finding provides a first indication that the human liver remains a young organ and constantly renews throughout a person’s lifetime.

As the organ comprises different cell types, we further separated nuclei from hepatocytes and nonhepatocytes. Genomic ^14^C concentrations suggest that hepatocytes are renewed over the whole human lifetime with a renewal rate of 20% per year and an average age of 2.5 years. Nonhepatocytes are only a small fraction of liver cells but represent a very diverse group with various functions. The largest populations include endothelial, Kupffer and stellate cells. However, a real characterization of their unique renewal dynamics is missing. Our obtained ^14^C data provide general insight into the turnover behavior of nonhepatocytes. We determined an average renewal rate of 20% per year, which is comparable to the turnover in hepatocytes. However, as we did not focus on characterizing the renewal rates of individual subpopulations, our ^14^C measurements provide only an estimate of the average turnover rate of all nonhepatocytes.

### Heterogeneity in hepatocyte renewal

Recent reports have questioned whether all hepatocytes contribute equally to tissue maintenance (Bangru and Kalsotra, 2020). Thus, we asked if there might be distinct, independent subpopulations with different turnover behaviors. As the liver is usually described as a quiescent organ (Berasain and Avila, 2015), we first tested whether there is indeed a pool of hepatocytes that do not participate in cellular renewal but remain quiescent over the lifetime (POP1q scenario, Fig. S4A). However, our data showed that at least 96% of all hepatocytes divide lifelong, providing no indication that a significant quiescent subpopulation of hepatocytes exists in the human liver. Accordingly, markers of permanent cell cycle arrest are physiologically not detectable in parenchymal liver cells (Aravinthan et al., 2013; Wang et al., 2018). Together, these observations suggest that virtually all hepatocytes participate in the organ’s renewal over a life span.

However, studies have repeatedly proposed that certain hepatocytes might possess a higher proliferative capacity than others. This possibility has been particularly discussed in the context of metabolic zonation (Manco et al., 2018). Therefore, we next questioned whether two independent hepatocyte pools exist in the liver, both contributing to organ renewal at different rates (scenario POP2, Fig. S4A). Our data show that these subpopulations have very similar turnover potential, challenging the idea of a more proliferative hepatocyte subpopulation.

Furthermore, these two scenarios testing different possibilities of heterogeneity in hepatocyte renewal show no considerably better fit of our ^14^C data compared with the simple POP1 scenario (Fig. S4A). Taken together, our data provide no evidence for the existence of independent hepatocyte subpopulations that contribute differently to liver renewal.

### Hepatocyte ploidy determines cellular renewal capacity

Polyploidy has been described in several organs of the body, including the heart (Bergmann et al., 2015) and the liver (Bou-Nader et al., 2019). However, the nature and biological role of ploidy is only poorly understood (Derks and Bergmann, 2020). In contrast to other organs, polyploidy in the liver does not contradict proliferation or regeneration in animal studies (Lin et al., 2020; Zhang et al., 2018). In humans, there is little knowledge on how polyploid cells, which represent a much lower fraction than in the murine organ, contribute to tissue renewal.

We show that human polyploid hepatocytes maintain their ability to renew. However, the chance to be renewed is more than seven-fold lower than that of diploid cells. Polyploid hepatocytes can remain in the liver for more than a decade, forming a subpopulation of long-lived cells. This apparent proliferative advantage of diploid over polyploid hepatocytes is supported by numerous publications (Chen et al., 2020; Wang et al., 2015; Wilkinson et al., 2019). However, in contrast to humans, most murine hepatocytes (90%) become polyploid starting approximately at the time of weaning (Duncan et al., 2010; Wang et al., 2014). A recent mouse study labeling all preexisting cells with ^15^N showed that 95% of all hepatocytes are not replaced within one and a half years (Arrojo e Drigo et al., 2019), indicating that most hepatocytes are long-lived. Whether the described small fraction of renewing murine hepatocytes might correspond to diploid hepatocytes with a higher turnover rate, as we report in humans, remains an open question and requires further investigation.

In mice, the distribution of polyploidy is strongly related to zonal localization (Matsumoto et al., 2020; Morales-Navarrete et al., 2015; Tanami et al., 2017). Therefore, it seems possible that different renewal capacities of the ploidy classes also reflect hepatocyte zonation. However, in the human liver, no particular ploidy distribution regarding liver zonation has been reported (Bou-Nader et al., 2019). This difference underlines that the age distribution of hepatocytes varies across species and that cell renewal in the mouse liver is different from that in humans.

### The polyploid hepatocyte population is mainly maintained by self-renewal

The increase in hepatocyte ploidy over the lifetime was described several decades ago (Kudryavtsev et al., 1993). However, there are few data on the mechanism of this ploidy change in the human liver. Aberrant cell cycle activity of diploid hepatocytes has been seen as the main driver of polyploidy in the liver (Celton-Morizur et al., 2009; Margall-Ducos et al., 2007). Therefore, we were interested in understanding how often aberrant cell cycle activity results in a change in ploidy class in humans. We observed that, independent of subject age, almost all diploid hepatocytes showed regular cell divisions (> 99%). At the same time, approximately 11% of polyploid cells entering the cell cycle return to diploidy. The fractions of human hepatocytes changing the ploidy level are in the same range as in murine models (Duncan et al., 2010).

We observed that an increase in the ploidy level is necessary to explain the initial generation of polyploid liver cells. However, the significant difference between the ^14^C concentrations in diploid and polyploid hepatocyte nuclei (Fig. 3F) already indicates that such ploidy transition processes play only a minor role. The more refined analysis with our mathematical model confirms that the exchange between the diploid and polyploid population is several-fold lower than the individual renewal rates of diploid and polyploid hepatocytes (birth and death of cells). We report that in young individuals, almost one-fifth of polyploid hepatocytes originate from diploid hepatocytes, while this fraction drops to less than 4% in older individuals (Fig. 5A). Thus, the majority of polyploid cells do not arise by a ploidy increase from diploid cells but due to proliferation of already existing polyploid hepatocytes. In the last decade, the importance of a bidirectional exchange between different ploidy levels for maintaining liver homeostasis has been emphasized in the ploidy conveyer model (Duncan et al., 2010), although a more recent publication suggests that the murine polyploid cell compartment remains stable over 18 months in homeostasis (Matsumoto et al., 2020). We observed that the contribution of polyploid cells to the diploid cell pool was less than 4% over the whole lifetime (Fig. 5B), playing only a minor role in cell generation. In conclusion, our model implies that the age-related increase in human hepatocyte polyploidy is mainly attributed to a net increase in newly generated cells within the individual polyploid hepatocyte pool.

### Continuous renewal of hepatocytes in the aging liver

It is important to understand the impact of aging on cell renewal in the human liver, as many chronic liver diseases show a higher prevalence in aged individuals (Kim et al., 2015). Until now, there have been only limited data available on age-dependent hepatocellular renewal (Macdonald, 1961) and its impact on liver function. Our model provides no evidence that the renewal rates of adult hepatocytes are affected by the subject’s age, with almost constant birth rates in diploid and polyploid hepatocytes, respectively (Fig. 4B). However, due to the relative increase in the size of the polyploid compartment in the aging liver, the average hepatocyte age and their age distribution change over the human life span (Fig. 4C-F).

Polyploid cells have been associated with aneuploidy and genomic instability (Storchova and Pellman, 2004). However, we found substantially lower turnover rates in polyploid hepatocytes, possibly being a protective mechanism against replicative senescence and somatic mutations. Moreover, polyploidy has been suggested to be an adaptive response forming a buffer against oxidative stress and DNA damage, increasing in liver aging and disease processes (Gentric and Desdouets, 2014).

In summary, ^14^C retrospective birth dating revealed that the human liver remains a young organ even in elderly individuals. Over the whole lifetime, we observed a constant and high turnover of hepatocytes that is strongly associated with ploidy level. Exchange between the ploidy classes plays only a minimal role in liver homeostasis in humans. Whether long-lived polyploid cells make the liver more susceptible to age-dependent diseases or act as a resilience factor to cope with cellular stress, thereby preventing loss of organ function and cancer, remains an important question to be addressed.

## Acknowledgments

We thank Magdalena Kretschmer and Irina Simonova for technical assistance and Steffen Rulands and Fabrizio Olmeda for discussions. This work was supported by the Flow Cytometry Facility, the Light Microscopy Facility and Histology Facility, all Core Facilities of the CMCB Technology Platform at TU Dresden. O.B. was supported by the Center for Regenerative Therapies Dresden, the Karolinska Institutet, the Swedish Research Council, the Ragnar Söderberg Foundation, and the Åke Wiberg Foundation. L.B. acknowledges support by the BMBF (grant 031L0033). The model simulations and Bayesian inference were performed on HPC resources granted by the ZIH at TU Dresden.

## Author Contributions

Conceptualization, P.H. and O.B.; Methodology, P.H., F.R., J.R., M.S., L.B. and O.B.; Software, F.R., J.R. and L.B.; Investigation, P.H., J.F. and O.B.; Resources, T.W., K.A., and H.D.; Writing – Original Draft, P.H., F.R. and O.B.; Writing – Review & Editing, P.H., F.R., L.B. and O.B.; Supervision, M.S., G.P., H.D., L.B. and O.B.; Funding Acquisition, L.B. and O.B.

## Figure titles and legends

**Supplementary figure S1: Flow cytometry and immunohistochemical analysis of liver cell nuclei**.

(A) HNF4α-based sorting strategy results in high purities of hepatocyte (98.3 ± 0.8%) and nonhepatocyte nuclear fractions (97.2 ± 1.5%) as confirmed by reanalyses. (B) Hepatocyte nuclei accounted for 73.7% of all liver nuclei, independent of subject age (n = 32, R = 0.06, p = 0.73). (C) Ki-67 expression in HNF4α-positive hepatocyte nuclei (n = 4) (median with interquartile range), scale bar 25 µm.

**Supplementary figure S2:** ^**14**^**C retrospective birth dating of human liver cells**.

(A and C) Genomic ^14^C concentrations of all liver nuclei (A) and nonhepatocyte nuclei (C) do not correlate with the atmospheric ^14^C concentration at the time of birth. (B and D) The estimated carbon age of all liver nuclei (B, n = 23, R = 0.02, p = 0.92) and nonhepatocyte nuclei (D, n = 11, R = 0.04, p = 0.91) is independent of the individual’s age and proves a continuous lifelong cellular turnover.

**Supplementary figure S3: Quantitative imaging and FACS-based strategies to determine the nuclear ploidy of hepatocytes**.

(A) Fluorescence intensity of genomic DNA and nuclear volume of hepatocyte nuclei identify different ploidy classes. (B) Observed ploidy profiles of hepatocyte nuclei are independent of the study approach (blue checker – determined by nuclear DRAQ5 fluorescence intensity with flow cytometry, and green checker – determined by nuclear size with quantitative imaging). (C) Comparison of quantitative imaging-based analysis of hepatocyte nuclei (squares) to previously published data by Kudryavtsev et al., 1993 (circles) confirms age-dependent changes in tetraploid (Gaussian regression, n = 316, R = 0.94) and octaploid cell populations (Gaussian regression, n = 316, R = 0.95). (D) Reanalyses of ploidy-sorted hepatocyte nuclei (HNF4α/DRAQ5) demonstrate a high level of sorting purity of diploid and polyploid hepatocyte nuclei fractions. (E) The ^14^C concentration of polyploid hepatocyte nuclei corresponds to a time point before the ^14^C formation date of diploid hepatocyte nuclei.

**Supplementary figure S4: Mathematical modeling predicts the dynamics of hepatocyte renewal**.

(A) Model sketches of the tested mathematical scenarios POP1, POP1q, POP2 and POP2p (β = birth rate, δ = death rate, κ = exchange rate, f = cellular fraction). (B) Analysis of individual hepatocyte ^14^C measurements with the POP1 scenario provides no evidence for age-related changes in the hepatocyte birth rate (n = 25, R = 0.11, p = 0.61). (C) The predicted ^14^C levels according to scenario POP2p (crosses) correspond to the measured ^14^C concentrations (dots) (connected with dashed line) (see supplementary methods).

## Table titles and legends

**Supplementary table S1: Analyzed human subjects**.

Patient characteristics and information about liver organ.

**Supplementary table S2:** ^**14**^**C measurements of human liver cell nuclei**.

^14^C values and related data of analyzed samples.

## Methods

### Tissue collection

Nondiseased human liver tissues were obtained by KI Donatum after informed consent from the individual or next of kin from 29 autopsies between 2004 and 2017 from the Swedish National Department of Forensic Medicine and from six patients who underwent liver resection surgery in 2017 at the University Hospital Carl Gustav Carus Dresden (Table S1). Ethical permission for these studies was granted by the regional ethics review board in Stockholm (Dnr 2005/1029-31/2 and Dnr 2010/313-31), and the Ethikkommission an der TU Dresden (EK 365082016 and EK251062017). The liver tissues were frozen and stored at -80 °C until further analysis.

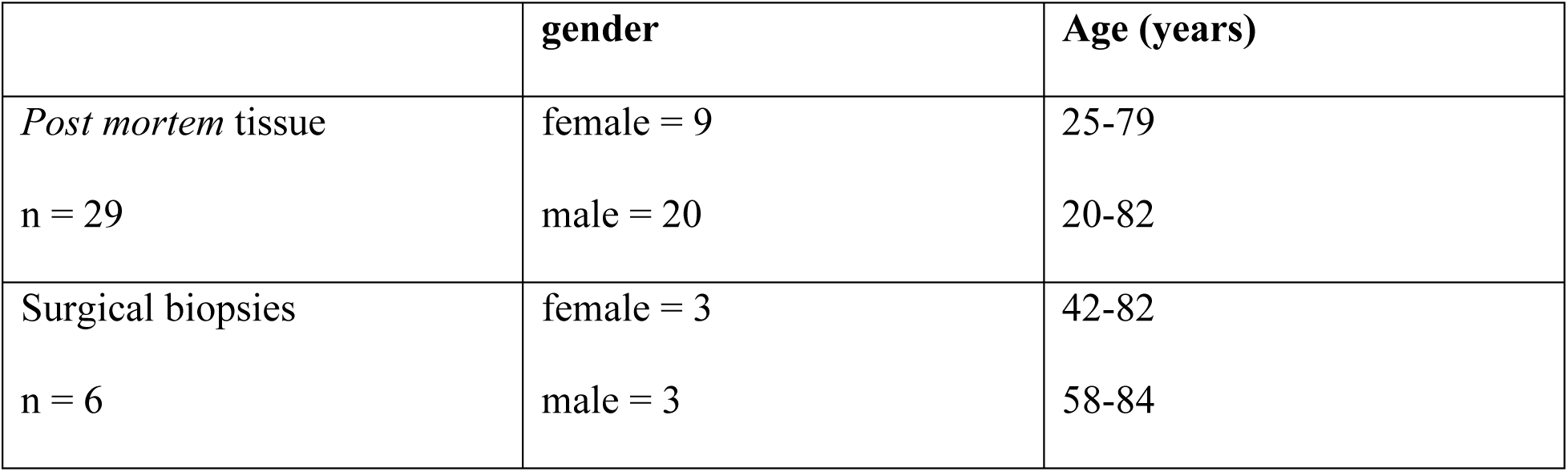

### Nucleus isolation

All steps were carried out at 4 °C. Approximately 4 g of liver tissue was homogenized in 200 ml of lysis buffer (0.32 M sucrose, 10 mM Tris-HCl, 5 mM CaCl_2_, 5 mM MgAc, 2 mM EDTA) using a domestic blender and dounce homogenizer. Samples were then suspended in sucrose solution (2.1 M sucrose, 10 mM Tris-HCl, 5 mM MgAc), layered onto cushions of pure sucrose solution and centrifuged for 1 h at 26,500 x*g*. The isolated nuclei were then resuspended in nuclear storage buffer (0.44 M sucrose, 10 mM Tris-HCl, 70 mM KCl, 10 mM MgCl_2_, 2 mM EDTA).

### Flow cytometry analysis and sorting

All steps were carried out at 4 °C. Isolated nuclei were stained overnight with HNF4α-antibody (Santa Cruz, sc-374229, RRID: AB_10989766) 1:1000, followed by 1 h incubation with Alexa Fluor-488 coupled secondary antibody (Jackson Immuno Research, 715-546-151, RRID: AB_2340850) 1:1000. DRAQ5 (Biostatus, DR51000) 1:5000 was added as nuclear stain directly before the analysis with a BD FACSAria™ II, III or Fusion. Single nuclei were identified and gated using FSC and SSC. Hepatocyte nuclei were identified and sorted based on their increased Alexa Fluor-488 fluorescence intensity. Separation of diploid and polyploid hepatocyte nuclei was based on DNA staining. Purities of the sorted populations were confirmed by reanalyses (Fig. S1A and Fig. S3D). Due to impurities, the measured ^14^C levels are actually a mixture of hepatocyte and nonhepatocyte ^14^C concentrations. For a paired measurement in hepatocytes and nonhepatocytes from the same individual, the purity-corrected ^14^C concentration was computed by solving the equations

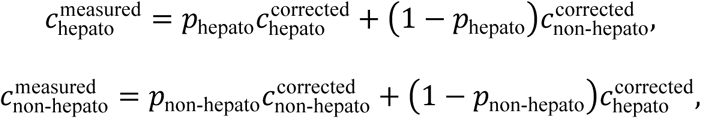

for the corrected ^14^C concentrations 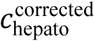 and 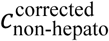. In the equations, *p*_hepato_ and *p*_non-hepato_ are the purities of the hepatocyte and the nonhepatocyte populations. Similarly, diploid and polyploid hepatocyte samples were corrected for purity for paired measurements. In this case, the above equations were changed to reflect that tetraploid cells contain twice as much DNA as diploid cells. The sorted liver cell nuclear pellets were frozen at -20 °C until further processing.

### DNA extraction

All steps were performed under clean room conditions to prevent any carbon contamination of the samples. Then, 500 µl DNA lysis buffer (1% SDS, 5 mM EDTA-Na2, 10 mM Tris-HCl, in UltraPure™ distilled water) and 6 µl proteinase K (20 mg/ml) were added to the samples and incubated overnight at 64 °C. Half the volume of 5 M NaCl was added. Then, the samples were agitated for 15 s and centrifuged for 3 min at 16000 x*g*. Supernatants were transferred into glass tubes, 3× 96% ethanol was added, and DNA was precipitated by inversion. Subsequently, DNA precipitates were washed three times in washing buffer (70% EtOH, 0.5 M NaCl, in UltraPure™ distilled water) and dissolved in 500 µl UltraPure™ distilled water. DNA quantity and purity were assessed with UV spectroscopy (NanoDrop).

### Accelerator mass spectrometry (AMS)

Purified DNA samples were analyzed blindly with AMS as described previously (Spalding et al., 2013). The DNA samples were lyophilized to dryness and reduced to graphite, and ^14^C/^12^C ratios were measured at the Department of Physics and Astronomy, Applied Nuclear Physics, Ion Physics, Uppsala University, Sweden, using a 170 kV commercial MICADAS accelerator mass spectrometer (IonPlus AG, Zurich, Switzerland). The carbon background was subtracted, as described previously (Salehpour et al., 2015; Spalding et al., 2013). All ^14^C data are reported as Δ^14^C, and the measurement error of each sample ranged between ± 8‰ and 43‰ (2 SD) Δ^14^C (Table S2).

### Mathematical modeling

See supplementary methods.

### Immunohistochemistry

Four percent paraformaldehyde-fixed human liver tissues were either paraffin embedded or frozen. Sections were cut at thicknesses of 6 µm (paraffin) and 40 µm (cryo). Paraffin sections were deparaffinized and rehydrated, and antigens were retrieved by 20 min incubation in boiling 0.05% citraconic anhydride buffer. The sections were incubated overnight with primary antibodies, followed by 1 h incubation with secondary antibodies. The HNF4α signal was amplified using a Tyramide Super Boost Kit (Life Technologies, B40933). Nuclei were counterstained with DRAQ5 (Biostatus, DR51000) 1:500. Images were collected with Apotome or LSM.

To assess the fraction of cycling hepatocytes in homeostasis, tissue sections were stained with primary antibodies against HNF4α/Prox1 and Ki-67. Images were collected with Apotome. Between 6,900 and 10,000 HNF4α/Prox1+ nuclei were manually evaluated for double positivity with Ki-67 using ImageJ software (version 1.52p).

Primary antibodies for IHC

| Antigen | Company | Catalog Number | Dilution | RRID |
| --- | --- | --- | --- | --- |
| Arginase-1 | Sigma Aldrich | HPA024006 | 1:1000 | AB_1844987 |
| HNF4α | Santa Cruz | sc-374229 | 1:500 | AB_10989766 |
| Ki-67 | Abcam | ab16667 | 1:1000 | AB_302459 |
| Ki-67 | Dako | M7240 | 1:100 | AB_2142367 |
| Prox1 | Abcam | ab11941 | 1:800 | AB_298722 |
| Slc2a2 | Atlas Antibodies | HPA028997 | 1:1000 | AB_2672860 |

Secondary antibodies for IHC

| **Antigen** | **Conjugate** | **Company** | **Catalog**<br>**Number** | **Dilution** | **RRID** |
| --- | --- | --- | --- | --- | --- |
| Anti-Rabbit<br>IgG (H+L) | Alexa Fluor®<br>555 | Abcam | ab150062 | 1:100 | AB_2801638 |
| Anti-Mouse<br>IgG (H+L) | Biotin-SP<br>(long spacer) | Jackson<br>Immuno<br>Research | 715-066-151 | 1:250 | AB_2340788 |
| Anti-Rabbit<br>IgG (H+L) | Alexa Fluor®<br>488 | Jackson<br>Immuno<br>Research | 711-546-152 | 1:250 | AB_2340619 |

### Analysis of nuclear ploidy and multinucleation

Immunohistochemistry was performed on 40 µm thick human liver sections. First, we visualized all nuclei (DRAQ5) and hepatocyte nuclei specifically (HNF4α) (n = 4). An LSM confocal microscope (63x oil objective) was used to take image stacks with z spacing 0.5 µm. The nuclei were automatically analyzed for their fluorescence intensity and size using ImageJ software (version 1.52p), confirming a correlation of these two characteristics.

For integrated analysis of nuclear and cellular ploidy, the LSM confocal microscope was used in conjunction with Visiomorph software. Tissue samples were stained for nuclei (DRAQ5) and hepatocyte membranes (Slc2a2) (n = 6). Hepatocyte nuclei were identified by their shape and localization in relation to Slc2a2-positive cell membranes. An automated meandering algorithm was applied to randomly choose more fields of view until a total of approximately 100 hepatocytes had been studied. All hepatocytes in one field of view were analyzed, and complete cell bodies were present in the tissue section. Multinucleation was visually determined utilizing Slc2a2 membrane staining by zooming through the optical layers of the section. Furthermore, hepatocyte nucleus diameters were measured at the maximal cross-sectional area.

To assign the nuclei to different ploidy classes based on their size (Bou-Nader et al., 2019; Tanami et al., 2017; Watanabe and Tanaka, 1982), a cluster analysis was performed for each individual case. The nuclear volumes were normalized to the modal values, and cut-offs were set to 1.5x (2n <> 4n), 3x (4n <> 8n) and 6x mode (8n <> 16n).

