## Supplemental Table 1 for "Diploid hepatocytes drive physiological liver renewal in adult humans"

**Supplementary table S1: Analyzed human subjects.**

Patient characteristics and information about liver organ.

| **Case** | **Year of birth** | **Age (years)** | **Gender** | **BMI** | **Liver weight (g)** | **Hepatic steatosis** | **Alcohol abuse** | **Collection type** | **Cause of death** | **Indication for surgery** |
| --- | --- | --- | --- | --- | --- | --- | --- | --- | --- | --- |
| ND022 | 1946 | 58 | M | 24.93 | 1955 | no | no | *Post mortem* tissue | suicide |  |
| ND023 | 1945 | 59 | M | 29.73 | 2653 | no | no | *Post mortem* tissue | cardiac infarction |  |
| ND252 | 1975 | 37 | M | 20.30 | 2013 | no | no | *Post mortem* tissue | cardiac infarction |  |
| ND253 | 1978 | 33 | F | 19.23 | 1322 | no | no | *Post mortem* tissue | motor car accident |  |
| ND255 | 1961 | 50 | F | 21.83 | 1454 | no | no | *Post mortem* tissue | suicide |  |
| ND256 | 1982 | 30 | M | 19.87 | 1672 | no | no | *Post mortem* tissue | suicide |  |
| ND257 | 1987 | 25 | M | 22.20 | 2041 | no | no | *Post mortem* tissue | intoxication |  |
| ND258 | 1968 | 43 | M | 30.99 | 1965 | slight | no | *Post mortem* tissue | intoxication |  |
| ND375 | 1992 | 23 | M | 28.93 | 2136 | no | no | *Post mortem* tissue | cardiac infarction |  |
| ND380 | 1942 | 73 | F | 27.22 | 2610 | no | no | *Post mortem* tissue | acute myocarditis |  |
| ND381 | 1985 | 31 | M | 22.55 | 1932 | no | no | *Post mortem* tissue | suicide |  |
| ND382 | 1982 | 34 | F | 24.84 | 2087 | no | no | *Post mortem* tissue | intoxication |  |
| ND383 | 1937 | 79 | F | 25.39 | 1590 | slight | no | *Post mortem* tissue | cardiac infarction |  |
| ND387 | 1995 | 21 | M | 23.51 | 1343 | no | no | *Post mortem* tissue | CO intoxication |  |
| ND388 | 1991 | 25 | M | 23.36 | 1716 | no | no | *Post mortem* tissue | unknown |  |
| ND390 | 1963 | 52 | M | 30.84 | 2654 | slight | no | *Post mortem* tissue | cardiac infarction |  |
| ND393 | 1943 | 73 | F | 41.93 | 1998 | no | no | *Post mortem* tissue | suicide |  |
| ND395 | 1971 | 46 | F | 23.42 | 1180 | no | no | *Post mortem* tissue | suicide |  |
| ND399 | 1997 | 20 | M | 18.39 | 1110 | no | no | *Post mortem* tissue | motor car accident |  |
| ND401 | 1979 | 37 | M | 29.30 | 2312 | no | no | *Post mortem* tissue | acute alcohol intoxication |  |
| ND402 | 1952 | 64 | F | 21.88 | 1520 | no | no | *Post mortem* tissue | suicide |  |
| ND403 | 1992 | 25 | F | 23.03 | 1512 | no | no | *Post mortem* tissue | suicide |  |
| ND404 | 1961 | 55 | M | 27.14 | 2198 | no | no | *Post mortem* tissue | unknown |  |
| ND405 | 1951 | 65 | M | 32.41 | 2824 | no | no | *Post mortem* tissue | acute alcohol intoxication |  |
| ND406 | 1983 | 33 | M | 21.28 | 1608 | no | no | *Post mortem* tissue | acute intoxication |  |
| ND407 | 1948 | 69 | M | 26.45 | 1552 | no | no | *Post mortem* tissue | suicide |  |
| ND408 | 1975 | 42 | M | 23.45 | 1692 | slight | no | *Post mortem* tissue | suicide |  |
| ND413 | 1963 | 54 | M | 23.30 | 1050 | no | no | *Post mortem* tissue | suicide |  |
| ND426 | 1935 | 82 | M | 24.28 | 1102 | no | no | *Post mortem* tissue | internal bleeding |  |
| HL#10 | 1932 | 84 | M | 22.65 | unknown | no | unknown | Surgical biopsy |  | intrahepatic cholangiocarcinoma |
| HL#11 | 1975 | 42 | F | 20.45 | unknown | no | unknown | Surgical biopsy |  | liver metastasis ovarial cancer |
| HL#12 | 1947 | 70 | F | 34.85 | unknown | slight | unknown | Surgical biopsy |  | liver metastasis breast cancer |
| HL#5 | 1952 | 64 | M | 27.41 | unknown | slight | unknown | Surgical biopsy |  | liver metastasis colon adenocarcinoma |
| HL#7 | 1935 | 82 | F | 18.73 | unknown | slight | unknown | Surgical biopsy |  | liver metastasis rectal carcinoma |
| HL#8 | 1959 | 58 | M | 19.37 | unknown | slight | unknown | Surgical biopsy |  | liver metastasis renal cell carcinoma |
