## Supplemental Table 2 for "Diploid hepatocytes drive physiological liver renewal in adult humans"

**Supplementary table S2: ^14^C measurements of human liver cell nuclei.**

^14^C values and related data of analyzed samples.

| **Case** | **Population** | **Δ^14^C** | **Error (± 2SD)** | **FACS purity (%)** | **Corrected Δ^14^C^a^** |
| --- | --- | --- | --- | --- | --- |
| ND022 | HNF4α+ | 75.0 | 9.4 | 99 |  |
| ND022 | HNF4α+ diploid | 78.6 | 17.2 | 94.8 |  |
| ND022 | unsorted | 77.2 | 11.6 |  |  |
| ND023 | HNF4α+ | 90.4 | 8.8 | 98 |  |
| ND023 | HNF4α+ diploid | 94.2 | 10.4 | 98.6 | 94.1 |
| ND023 | HNF4α+ polyploid | 105.4 | 11.6 | 93.1 | 106.3 |
| ND023 | unsorted | 81.2 | 8.8 |  |  |
| ND252 | HNF4α+ | 88.7 | 12.3 | 97 |  |
| ND252 | HNF4α+ diploid | 86.8 | 11.08 | 96.2 | 86.4 |
| ND252 | HNF4α+ polyploid | 104.91 | 12.4 | 94.0 | 106.0 |
| ND253 | HNF4α+ | 55.0 | 8.0 | 99 |  |
| ND253 | HNF4α+ diploid | 73.7 | 4.6 | 96.5 |  |
| ND255 | HNF4α+ | 89.0 | 11.9 | 98 |  |
| ND255 | HNF4α+ diploid | 80.5 | 15.7 | 88.7 | 78.5 |
| ND255 | HNF4α+ polyploid | 113.0 | 12.9 | 98.6 | 113.2 |
| ND256 | HNF4α+ | 5.8 | 19.6 | 98 |  |
| ND257 | HNF4α+ | 27.9 | 22.4 | 95 |  |
| ND258 | HNF4α+ | 71.9 | 9.9 | 99 | 72.0 |
| ND258 | HNF4α- | 58.6 | 14.9 | 99 | 58.5 |
| ND375 | HNF4α+ | 3.7 | 32.0 | 99 |  |
| ND380 | HNF4α+ | 67.5 | 14 | 98 |  |
| ND380 | unsorted | 78.4 | 8.0 |  |  |
| ND381 | unsorted | 31.5 | 8.0 |  |  |
| ND382 | HNF4α+ | 32.8 | 15.0 | 98 |  |
| ND382 | unsorted | 29.2 | 10.0 |  |  |
| ND383 | HNF4α+ diploid | 51.9 | 9.6 | 93 | 52.1 |
| ND383 | HNF4α+ diploid | 41.2 | 10.5 | 93 | 41.0 |
| ND383 | HNF4α+ polyploid | 46.9 | 10.7 | 92.8 | 46.9 |
| ND383 | HNF4α- | 37.7 | 13.9 | 93 |  |
| ND383 | unsorted | 51.4 | 14.3 |  |  |
| ND387 | HNF4α+ | 26.7 | 23.3 | 99 | 26.8 |
| ND387 | HNF4α- | 15.0 | 43.1 | 98 | 14.8 |
| ND387 | unsorted | 59.3 | 27.8 |  |  |
| ND388 | unsorted | 25.3 | 7.8 |  |  |
| ND390 | HNF4α+ | 28.0 | 19.0 | 99 |  |
| ND390 | unsorted | 41.2 | 13.8 |  |  |
| ND393 | HNF4α+ | 75.9 | 17 | 98 |  |
| ND393 | unsorted | 65.9 | 9.9 |  |  |
| ND395 | HNF4α+ | 47.8 | 19.9 | 98 | 47.4 |
| ND395 | HNF4α- | 67.0 | 38.6 | 97 | 67.7 |
| ND395 | unsorted | 78.0 | 12.5 |  |  |
| ND399 | HNF4α+ | 31.7 | 8.1 | 99 |  |
| ND399 | unsorted | 35.3 | 11.7 |  |  |
| ND401 | HNF4α+ | 52.4 | 18.0 | 98 | 51.8 |
| ND401 | HNF4α- | 85.2 | 21.2 | 97 | 86.3 |
| ND401 | unsorted | 43.5 | 20.0 |  |  |
| ND402 | HNF4α+ | 39.1 | 23.1 | 99 | 38.2 |
| ND402 | HNF4α- | 131.1 | 41.4 | 98 | 133.0 |
| ND402 | unsorted | 69.1 | 23.9 |  |  |
| ND403 | HNF4α+ | 46.6 | 9.5 | 98 | 46.7 |
| ND403 | HNF4α- | 44.5 | 18.0 | 97 | 44.5 |
| ND403 | unsorted | 54.1 | 9.8 |  |  |
| ND404 | HNF4α+ | 29.8 | 9.3 | 98 |  |
| ND404 | unsorted | 34.5 | 12.3 |  |  |
| ND405 | HNF4α+ | 21.6 | 8.4 | 99 | 21.0 |
| ND405 | HNF4α- | 86.7 | 19.6 | 98 | 88.1 |
| ND405 | unsorted | 26.1 | 10.7 |  |  |
| ND406 | HNF4α+ | 31.7 | 20.7 | 98 |  |
| ND407 | HNF4α+ | 41.3 | 9.3 | 99 |  |
| ND407 | unsorted | 41.5 | 8.0 |  |  |
| ND408 | HNF4α+ | 32.5 | 8.9 | 98 |  |
| ND408 | unsorted | 48.4 | 9.0 |  |  |
| ND413 | HNF4α+ | 18.8 | 13.2 | 99 |  |
| ND426 | HNF4α+ | 19.5 | 9.2 | 99 | 19.6 |
| ND426 | HNF4α- | 13.3 | 18.6 | 97 | 13.1 |
| HL#7 | HNF4α+ | 31.2 | 15.3 | 98 |  |
| HL#7 | unsorted | 25.6 | 15.7 |  |  |
| HL#8 | HNF4α+ | 7.4 | 8.7 | 98 | 6.6 |
| HL#8 | HNF4α- | 50.0 | 14.8 | 98 | 50.9 |
| HL#8 | unsorted | 26.3 | 8.9 |  |  |
| HL#10 | HNF4α- | 16.5 | 14.3 | 98 |  |
| HL#10 | unsorted | 13.0 | 9.2 |  |  |
| HL#11 | HNF4α+ diploid | 16.2 | 12.8 | 85.4 | 13.0 |
| HL#11 | HNF4α+ polyploid | 39.6 | 13.7 | 91 | 41.6 |
| HL#12 | HNF4α+ | 24.6 | 13.2 | 97 |  |
| HL#12 | unsorted | 19.2 | 13.1 |  |  |

^a^see supplementary methods
