## Supplementary figures and images for "Diploid hepatocytes drive physiological liver renewal in adult humans"

### Supplemental Figures

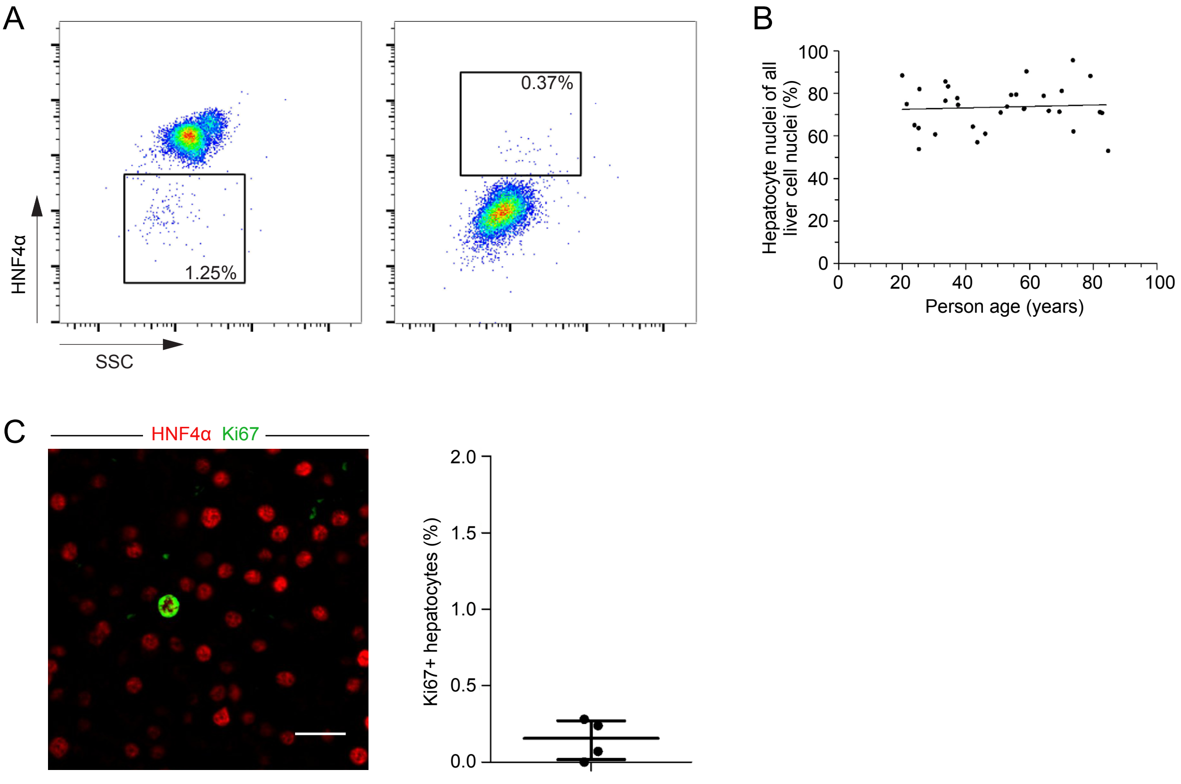


**Supplementary figure 1**

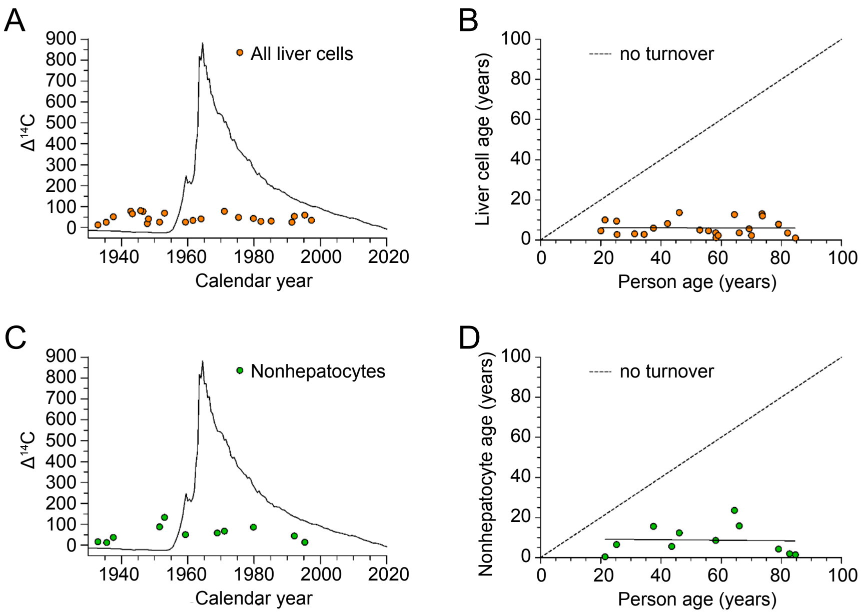


**Supplementary figure 2**

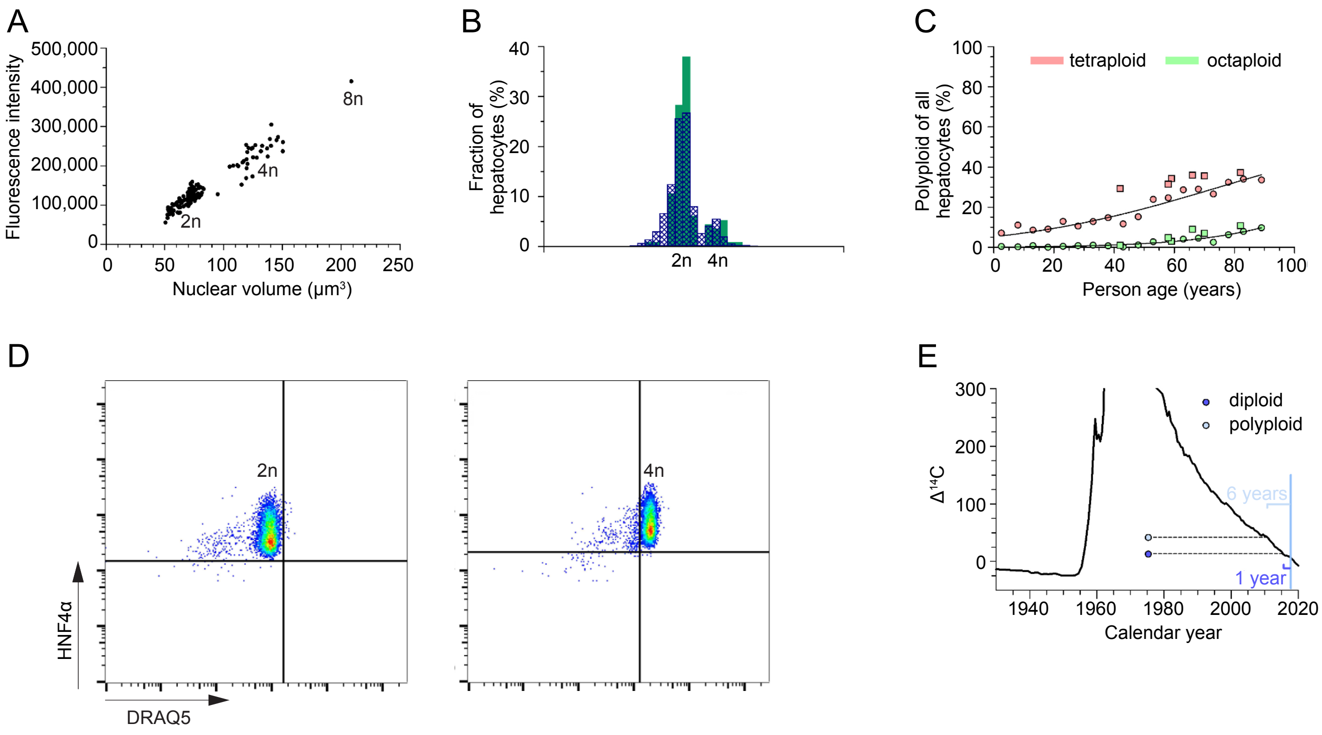


**Supplementary figure 3**

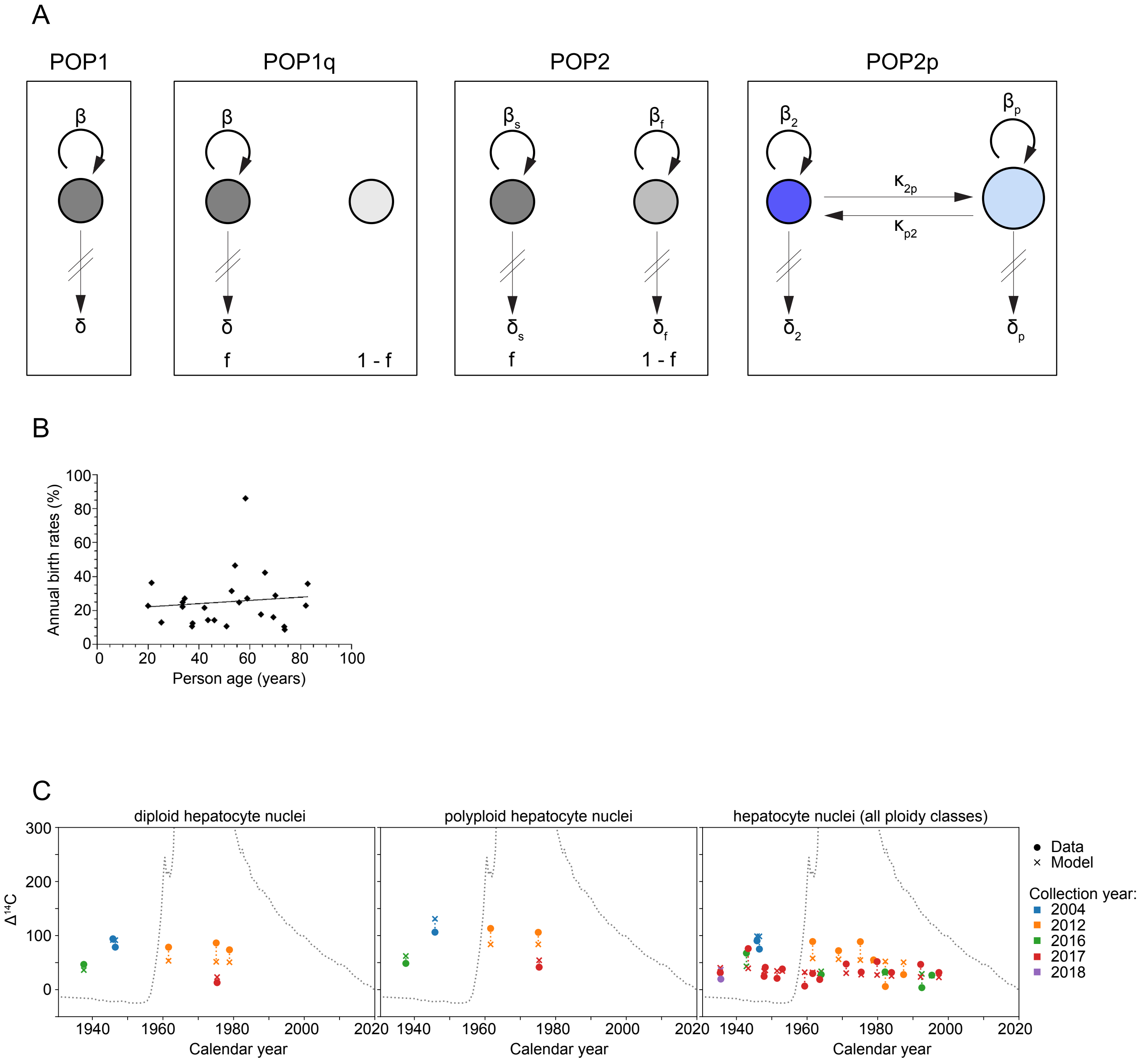


**Supplementary figure 4**
