## Supplemental Methods for "Diploid hepatocytes drive physiological liver renewal in adult humans"

**Supplementary methods**

In the following, we derive the mathematical models, describe the methods for parameter estimation and model selection, and present the results that are based on the models in detail.

**Mathematical model of $^{14}$C concentration dynamics**

Here, we develop a mathematical model that predicts the genomic $^{14}$C concentration dynamics for a self-renewing cell population. To date, label concentration dynamics have been modeled with age-, concentration- and division-structured cell populations (Bernard et al., 2010; Hasenauer et al., 2012; Hross and Hasenauer, 2016; Schittler et al., 2013). In particular, the dynamics of $^{14}$C concentrations were modeled with an age-structured cell population, assuming that mainly a rapidly cycling stem cell population contributes to tissue homeostasis (Réu et al., 2017; Spalding et al., 2013; Yeung et al., 2019). Instead, our model rests upon the assumption that hepatocytes are self-renewing cells, as demonstrated in several recent reports (Malato et al., 2011; Schaub et al., 2014; Wang et al., 2017; Yanger et al., 2014).

We describe the state of a cell population *i* with a density in the $^{14}$C concentration space, $n_i(c)$, where $n_i$ is the density of cells with $^{14}$C concentration *c*. Cells divide at rate $\beta_i(t)$ and die at rate $\delta_i(t)$. We model the cell density dynamics in a subject with the following population balance equation:

$$\frac{\partial n_i(c)}{\partial t} = \overbrace{2\beta_i(t) \int dc' \, f_i\big(c, c' \big| c_a(t+b)\big) \, n_i(c')}^{\text{cell birth by division}} - \overbrace{\big(\beta_i(t) + \delta_i(t)\big) \, n_i(c)}^{\text{cell loss by division and death}},$$

where $c_a(t+b)$ is the atmospheric $^{14}$C concentration adjusted for one year of delay along the food chain, *b* is the time of birth, and *t* is the age of the subject. The kernel $f_i\big(c, c' | c_a(t+b)\big)$ describes the fraction of dividing cells with $^{14}$C concentration $c'$ that become cells with $^{14}$C concentration c given an atmospheric $^{14}$C concentration of $c_a(t+b)$. We assume that new DNA, which is synthesized during DNA duplication in the cell cycle, has an atmospheric $^{14}$C concentration. Then, the mean $^{14}$C concentration of the two daughter cells is $\frac{c'+c_a(t+b)}{2}$. To fulfil this condition, the kernel must be symmetric around $\frac{c'+c_a(t+b)}{2}$, i.e.,

$$f_i\left(\frac{c' + c_a(t + b)}{2} - x, c'|c_a(t + b)\right) = f_i\left(\frac{c' + c_a(t + b)}{2} + x, c'|c_a(t + b)\right).$$

Furthermore, the kernel is cell number conserving, i.e.,

$$\int_0^\infty dc\, f_i(c, c'|c_a(t + b)) = 1.$$

In scenario POP1 (Fig. S4A), there is only one population of cells that divide and die at constant rates. Assuming the total cell number $N = \int_0^\infty dc\, n(c)$ is constant, we obtain $\beta = \delta$, i.e., the dynamics are described by a single turnover rate that equals the cell birth and the cell death rate. Now, we can derive the ordinary differential equation,

$$\frac{\partial \bar{c}}{\partial t} = \beta\, (c_a(t + b) - \bar{c}),$$

for the mean $^{14}$C concentration $\bar{c} = \int_0^\infty dc\, c\, n(c)$.

In scenario POP1q (Fig. S4A), there is a renewing population with turnover that is described by the equations governing scenario POP1 and a quiescent population without turnover. The relative size of the renewing population is $f$ ($0 \leq f \leq 1$). The predicted $^{14}$C concentration for the measurable population average is then given by $f\bar{c} + (1 - f)c_a(b)$.

In scenario POP2 (Fig. S4A), there are two renewing populations with independent slow and fast turnover rates $\beta_s$ and $\beta_f$, respectively. Each population is described by the equations governing scenario POP1. It is useful to parameterize the model with the difference in turnover rates $\Delta\beta = \beta_f - \beta_s$. The fraction of the population with slower turnover is given by $f$. The predicted $^{14}$C concentration as the measurable population average is then given by $f\bar{c}_s + (1 - f)\bar{c}_f$.

Furthermore, we define a more complex scenario POP2p (Fig. 4A and S4A) that accounts for the fact that hepatocytes exist in different states of ploidy. We model a diploid (2n) and a polyploid cell population (pn). As most polyploid hepatocytes are tetraploid (Fig. S3C), we neglected higher ploidy levels. We assume that binucleated diploid (2x 2n) and mononucleated tetraploid cells (4n) behave identically. In this scenario, cell turnover occurs due to (a) cell division at rates $\beta_2$ and $\beta_p$ for diploid and polyploid hepatocytes, respectively; (b) cell death at rates $\delta_2$ and $\delta_p$ for diploid and polyploid hepatocytes, respectively; and (c) exchange between the ploidy populations whereby diploid cells convert to polyploid cells by unfinished cell division at rate $\kappa_{2p}$, and conversely, polyploid cells convert to diploid cells at rate $\kappa_{p2}$. We derive the population balance equations

$$\frac{\partial n_2(c)}{\partial t} = \overbrace{2\beta_2(t)\int dc' f_2\big(c,c'|c_a(t+b)\big)\, n_2(c')}^{\text{cell birth by division}} + \overbrace{2\kappa_{p2} n_p(c)}^{\text{fission of pn cells}} - \overbrace{\big(\beta_2(t)+\delta_2+\kappa_{2p}\big)n_2(c)}^{\text{cell loss due to division, death and ploidy increase}}, \quad (1)$$

$$\frac{\partial n_p(c)}{\partial t} = \overbrace{2\beta_p(t)\int dc' f_p\big(c,c'|c_a(t+b)\big)\, n_p(c')}^{\text{cell birth by division}} + \overbrace{\kappa_{2p}\int dc'\, f_{2p}\big(c,c'|c_a(t+b)\big) n_2(c')}^{\text{unfinished division of 2n cells}} - \overbrace{\big(\beta_p(t)+\delta_p+\kappa_{p2}\big)n_p(c)}^{\text{cell loss due to division, death and ploidy decrease}}. \quad (2)$$

Unfinished cell divisions of 2n cells with $^{14}$C concentration c' always give rise to a polyploid cell with $^{14}$C concentration $\frac{c'+c_a(t)}{2}$ and hence

$$f_{2p}\big(c,c'|c_a(t+b)\big) = \delta\left(c - \frac{c' + c_a(t+b)}{2}\right),$$

where $\delta$ is the Dirac delta function. The total cell numbers are given by

$$N_{2/p} = \int_0^\infty dc\, n_{2/p}(c)$$

and hence with (1), (2):

$$\frac{dN_2}{dt} = \big(\beta_2(t) - \delta_2 - \kappa_{2p}\big)N_2 + 2\kappa_{p2}N_p, \quad (3)$$

$$\frac{dN_p}{dt} = \big(\beta_p(t) - \delta_p - \kappa_{p2}\big)N_p + \kappa_{2p}N_2. \quad (4)$$

We constrain this scenario by assuming that the total amount of DNA remains constant,

$$N_2 + 2N_p = const. \quad (5)$$

Furthermore, we constrain our model such that it agrees with previously measured ploidy dynamics (Kudryavtsev et al., 1993) (Fig. 3B). To do so, we fitted a univariate spline of second order p(t) to the ploidy data (positions of interior knots of the spline: 2.5, 2.5, 2.5, 89.0, 89.0, 89.0; spline coefficients: 0.918, 0.884, 0.542) and then required that

$$p(t) = \frac{N_2(t)}{N_2(t) + N_p(t)}. \tag{6}$$

The mean $^{14}C$ concentrations in the 2n and pn populations are given by

$$\bar{c}_{2/p} = \frac{\int dc\, c\, n_{2/p}(c)}{N_{2/p}}.$$

Using (1-6), we obtain

$$\frac{d\bar{c}_2}{dt} = \left(2\kappa_{p2}\frac{p(t)-1}{p(t)} - \beta_2(t)\right)\bar{c}_2 - 2\kappa_{p2}\frac{p(t)-1}{p(t)}\bar{c}_p + \beta_2(t)c_a(t+b), \tag{7}$$

$$\begin{aligned}\frac{d\bar{c}_p}{dt} &= -\frac{1}{2}\kappa_{2p}\frac{p(t)}{p(t)-1}\bar{c}_2 + \left(\kappa_{2p}\frac{p(t)}{p(t)-1} - \beta_p(t)\right)\bar{c}_p \\ &\quad + \left(\beta_p(t)\frac{p(t)+1}{p(t)-1} + \frac{1}{2}\kappa_{2p}\frac{p(t)}{p(t)-1}\right)c_a(t+b),\end{aligned} \tag{8}$$

$$\beta_2(t) = \delta_2 + \kappa_{2p} + 2\kappa_{p2}\frac{p(t)-1}{p(t)} - 2\frac{dp(t)}{dt}\frac{1}{(p(t)-2)p(t)}, \tag{9}$$

$$\beta_p(t) = \delta_p + \kappa_{p2} + 2\kappa_{2p}\frac{p(t)}{p(t)-1} - \frac{dp(t)}{dt}\frac{1}{2-3p(t)+p(t)^2}. \tag{10}$$

For the initial condition, we assume that all cells have the current atmospheric $^{14}C$ concentration at the time of subject birth, i.e., $\bar{c}_{2/p}(t=0) = c_a(b)$. This fully specifies the dynamics of the mean $^{14}C$ concentration. We would like to note that this scenario is too complex to describe cell turnover with a simple turnover rate. We need to specify the death rates and the exchange rates to fully define the model. The time-dependent cell birth rates can then be computed using the equations above. We implemented a numerical solution of our model in Python (https://github.com/fabianrost84/Heinke_2020).

**Parameter estimation and model selection**

Experimentally, we isolated hepatocyte nuclei and determined the mean $^{14}C$ concentrations for different mixtures of diploid and polyploid cell populations. Diploid nuclei (2n) can originate from mononucleated diploid (2n) and binucleated tetraploid cells (2x 2n), and polyploid nuclei (pn) can originate from mono- and binucleated polyploid hepatocytes. Nonploidy-sorted nuclei (2n/pn) originate from cells of all possible nucleation and ploidy levels. From scenario POP2p, we can predict the experimentally measured $^{14}C$ concentrations by weighting the $^{14}C$ concentrations $\bar{c}_2$ and $\bar{c}_p$ according to the relative fractions of ploidy levels in the hepatocyte populations:

$$\bar{c}(t) = \frac{w_2^i \bar{c}_2(t) + 2w_p^i \bar{c}_p(t)}{w_2^i + 2w_p^i},$$

where $w_{2/p}^i$ are the weights of the 2n/pn populations for a sort of type *i*, and the factor 2 reflects that pn cells have double the amount of DNA. The weights are given by

$$\begin{aligned} w_2^{2n} &= p(t), & w_p^{2n} &= p_{2\times 2n}(t), \\ w_2^{pn} &= 0, & w_p^{pn} &= 1, \\ w_2^{2n/pn} &= p(t), & w_p^{pn} &= 1 - p(t). \end{aligned}$$

Here, $p_{2\times 2n}(t)$ yields the relative fraction of binucleated tetraploid hepatocytes. For all other scenarios, we ignored the different compositions of the samples.

We used an additive Gaussian noise model to define a likelihood for our data. This noise has contributions from actual subject-to-subject variability and from measurement errors. We assumed that the variance $\sigma^2$ is constant for all samples. We estimated parameter ranges with Bayesian inference. Depending on the model, we used uniform priors in the log space for the unknown rate parameters:

$$\log_{10} \delta_i, \log_{10} \Delta\beta, \log_{10} \beta_s \sim \mathcal{U}(\log_{10} 10^{-6}\text{years}, \log_{10} 10^{1}\text{years})$$

$$\log_{10} \kappa_{2p}, \log_{10} \kappa_{p2} \sim \mathcal{U}(\log_{10} 10^{-3}\text{years}, \log_{10} 10^{1}\text{years}),$$

and a uniform prior for $\sigma$ and f:

$$\sigma \sim \mathcal{U}(0, 0.2),$$

$$f \sim \mathcal{U}(0, 1)$$

We eliminated the division rates in scenario POP2p and the death rates in the other scenario due to the cell number constraints. Mathematically, these rates can become negative. However, because a negative rate is physically not possible, an infinitely unlikely value must be defined for the likelihood. For such cases, the likelihood is set to $-\infty$.

First, we calculated the maximum a posteriori probability estimate with the Broyden–Fletcher–Goldfarb–Shanno optimizer using 100 randomly drawn initial conditions (Fröhlich et al., 2017; Hasenauer et al., 2012; Virtanen et al., 2020). To numerically solve the Bayesian inference problem, we used Markov chain Monte Carlo (MCMC) sampling as implemented in emcee: The MCMC Hammer (Foreman-Mackey et al., 2013). For each scenario, we used 3000 samples, of which 1000 were used for the burn-in phase, and the number of chains was 50 times the number of parameters. The initial values for the unknown parameters were drawn from a multivariate normal distribution in log space for all parameters except sigma. This normal distribution was centered around the maximum a posteriori probability estimate and has a standard deviation of 0.1.

To select the model with the highest predictive power, we employed leave-one-out cross validation (LOO). We estimated LOO from the MCMC chains using Pareto-smoothed importance sampling (Vehtari et al., 2017).

### Bayesian inference results

MCMC sampling resulted in empirical distributions that approximate the posterior distributions of the estimated parameters. These posterior distributions are visualized in Fig. A-D for the four hepatocyte scenarios, in Fig. E for nonhepatocytes, and Fig. F for nonsorted liver nuclei. From these posterior distributions, we estimated parameter values and confidence intervals (Table A).

### Hepatocytes

When using the simplest scenario POP1, we estimated a turnover rate of 20%/year (CI: [16%/year, 23%/year) (Table A, Fig. A).

When we added a quiescent population in scenario POP1q, we found a similar turnover rate of 25%/year, (CI: [16%/year, 31%/year]) (Table A, Fig. B). We estimated a 97% (CI: [96%, 100%]) fraction of renewing cells, which indicates that the vast majority of hepatocytes are renewed. Furthermore, the added model complexity did not result in a higher predictive power, as indicated by the decreased LOO information criterion.

Next, we fitted scenario POP2 to test for the existence of two hepatocyte populations with different turnover rates (Fig. C). For the turnover rate of the slower population, we estimated 16%/year (CI: [3%/year, 22%/year]), which is again similar to the turnover rates estimated for the two scenarios above. We found that the slow cycling population would contain 40% (CI: [0%, 58%]) of the hepatocytes. The faster cycling population turns over with a rate 6% higher than the slower population (CI: [0%/year, 44%/year]). Hence, this scenario does not exclude that the two populations actually turn over with the same rate or with very similar rates. The added complexity of the two cell populations only led to a marginal increase in predictive power compared with scenario POP1 (Table A). Taken together, we do not find evidence of two independent hepatocyte populations with clearly distinct turnover rates.

In contrast, the most complex scenario POP2p, which explicitly incorporates polyploidization of the hepatocytes, produced the best fit of the data with a model weight of 49% (Table A, Fig. D). This scenario predicts clear differences in turnover for diploid and polyploid hepatocytes with cell death rates of 80%/year (CI: [0%/year, 156%/year]) and 8%/year (CI: [4%/year, 12%/year]), respectively. Remarkably, the exchange rates between the diploid and the polyploid population are comparably low. We found 0.3%/year (CI: [0.1%/year, 0.4%/year] for diploid to polyploid and 1%/year (CI: [0%/year, 3%/year]) for polyploid to diploid. Using equations (9) and (10), we can compute the time-dependent birth rates (Fig. 4B). Both rates remain nearly constant. The diploid population divides much faster ($\beta_2 = 77\%/\text{year}$ for a 50-year-old subject) than the polyploid population ($\beta_p = 11\%/\text{year}$ for a 50-year-old subject).

We used the best-fitting scenario to estimate the number of diploid and polyploid hepatocytes that are born every day. For a short time interval, $t_0$, these numbers can be estimated as

$$n_2(t) = \beta_2 p(t) n_0 t_0,$$

$$n_p(t) = \beta_p \big(1 - p(t)\big) n_0 t_0,$$

where $n_0$ is the total number of hepatocytes in the liver. Assuming there are approximately $10^{11}$ hepatocytes in an average human liver (Nagy et al., 2020), we estimate that approximately 167 million diploid and 6 million polyploid hepatocytes are born every day in a 50-year-old adult human liver.

### Nonhepatocytes and nonsorted liver nuclei

For the nonhepatocyte sorts and the nonsorted liver nuclei, we used scenario POP1 to estimate an average turnover rate. We found an average turnover rate of 20%/year (CI: [5%/year, 44%/year]) and 19%/year (CI: [14%/year, 23%/year]) for the nonhepatocyte sorts and nonsorted liver nuclei, respectively.

### Single-cell-based model to determine cell age distribution

We simulated a single-cell-based model to compute the cell age distribution for scenario POP2p (Fig. 4C-F). The state of the model at time $t$ is given by the birth times $a_i$ ($i$=1…n) and $b_i$ ($j$=1…m) of $n$ diploid cells (2n) and $m$ polyploid cells (pn), respectively. We simulate the dynamics with a fixed time step $\Delta t$. At each time step, the age of all cells is increased by $\Delta t$. Furthermore, the following events may occur for each cell at every time step: (a) 2n and pn cells divide with probability $\beta_2(t) \times \Delta t$ and $\beta_p(t) \times \Delta t$, respectively. If cell division occurs, the corresponding cell is removed, and two new cells with an age of 0 are added. (b) 2n and pn cells die with probability $\delta_2(t) \times \Delta t$ and $\delta_p(t) \times \Delta t$, respectively. If cell death occurs, the corresponding cell is removed. (c) A 2n cell turns into a pn cell with the probability $\kappa_{2p} \times \Delta t$, and a pn cell turns into two 2n cells with probability $\kappa_{p2} \times \Delta t$. In both cases, the age of new cells is set to 0. If multiple events are selected for a single cell in a single time step, then only one of them is randomly selected with probabilities that are proportional to the corresponding rates. For a small time step, $\Delta t$, the cell number dynamics of the single-cell-based model converge to the solution of equations (3) and (4). Additionally, the birth times of the cells allow the cell age distribution to be calculated. We used the best-fit parameters for scenario POP2p (Table A) to parameterize the model. The simulation was initialized with $10^7\ p(0)$ 2n cells and $10^7(1 - p(0))$ pn cells. We performed the model until $t$ = 80 years using a time step of $\Delta \mathrm{t} = 10^{-3}$ years.

**Table A: Model selection and parameter estimates for the different model scenarios that were fitted to the hepatocyte data.**

LOO: value of the leave-one-out information criterion; lower values indicate higher predictive power (best model). Weights indicate the probability of each scenario being the best model among the tested scenarios.

| Scenario | LOO | Weight | Parameter | Value | 1-σ Confidence Interval |
|---|---|---|---|---|---|
| **POP1** | 21.6 | 18% | β | 20%/year | [16%/year - 23%/year] |
| | | | σ | 33‰ | [27‰ - 38‰] |
| **POP1q** | 21.1 | 12% | β | 25%/year | [16%/year - 31%/year] |
| | | | f | 97% | [96% - 100%] |
| | | | σ | 34‰ | [27‰ - 39‰] |
| **POP2** | 21.9 | 22% | $\beta_s$ | 16%/year | [3%/year - 22%/year] |
| | | | Δβ | 6%/year | [0%/year - 44%/year] |
| | | | f | 37% | [0% - 58%] |
| | | | σ | 33‰ | [27‰ - 38‰] |
| **POP2p** | 22.6 | 49% | $\kappa_{2p}$ | 0.3%/year | [0.1%/year - 0.4%/year] |
| | | | $\kappa_{p2}$ | 1%/year | [0%/year - 3%/year] |
| | | | $\delta_2$ | 80%/year | [0%/year - 156%/year] |
| | | | $\delta_p$ | 8%/year | [4%/year - 12%/year] |
| | | | σ | 31‰ | [25‰ - 36‰] |

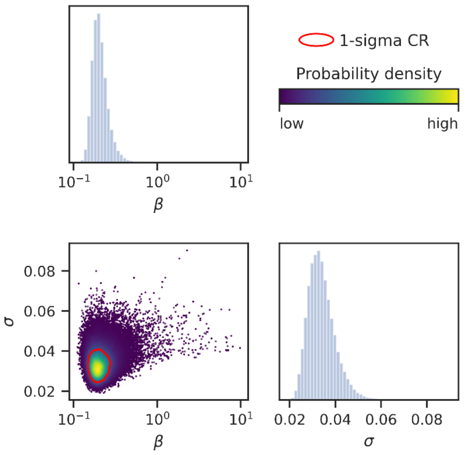


**Figure A: Posterior distributions of hepatocyte turnover parameters for the POP1 scenario estimated by MCMC sampling.** On diagonal panels: Marginal distributions of the model parameters. Off-diagonal panels: Marginal distributions of parameter pairs. Each dot is a sample from the MCMC chain, and the dots are color coded according to the local probability density. The 1-σ confidence region (CR) is enclosed by the red contour and contains the true parameter with 68% probability.

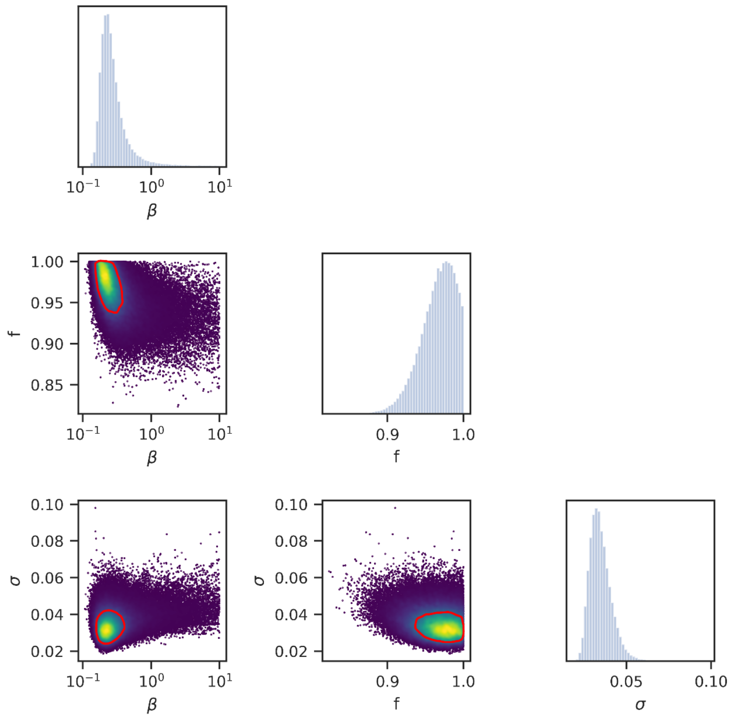


**Figure B: Posterior distributions of hepatocyte turnover parameters for the POP1q scenario estimated by MCMC sampling.**

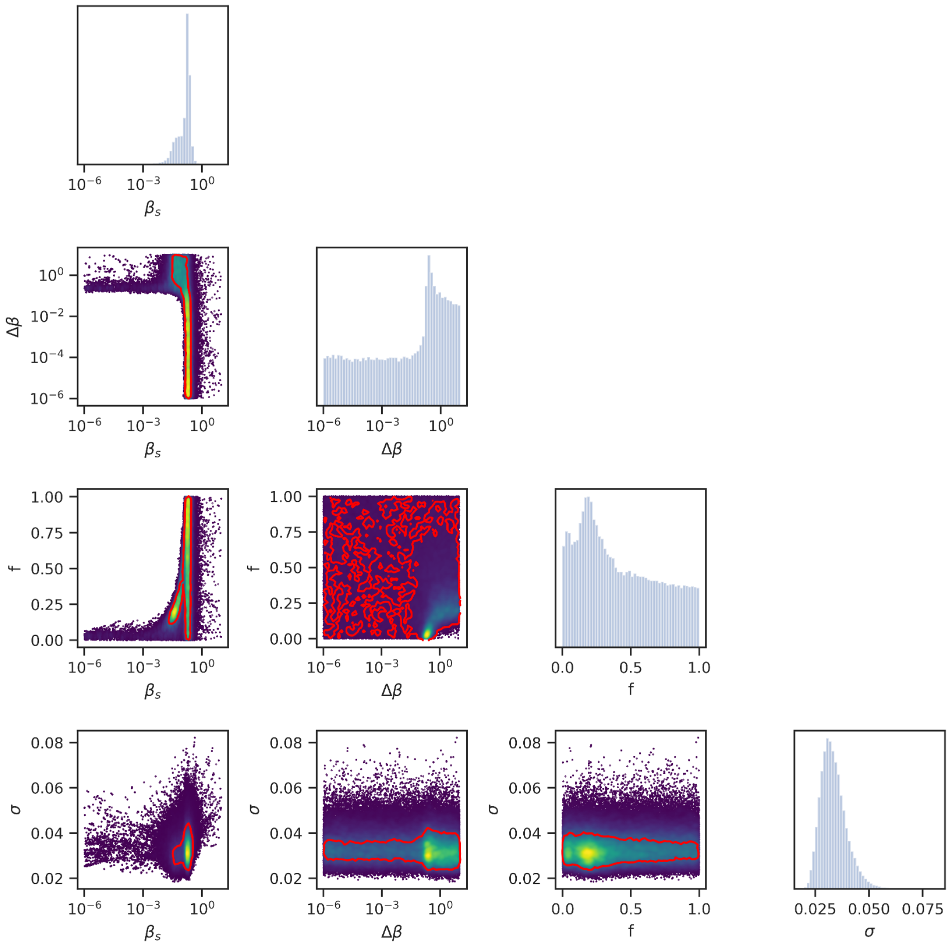


**Figure C: Posterior distributions of hepatocyte turnover parameters for the POP2 scenario estimated by MCMC sampling.**

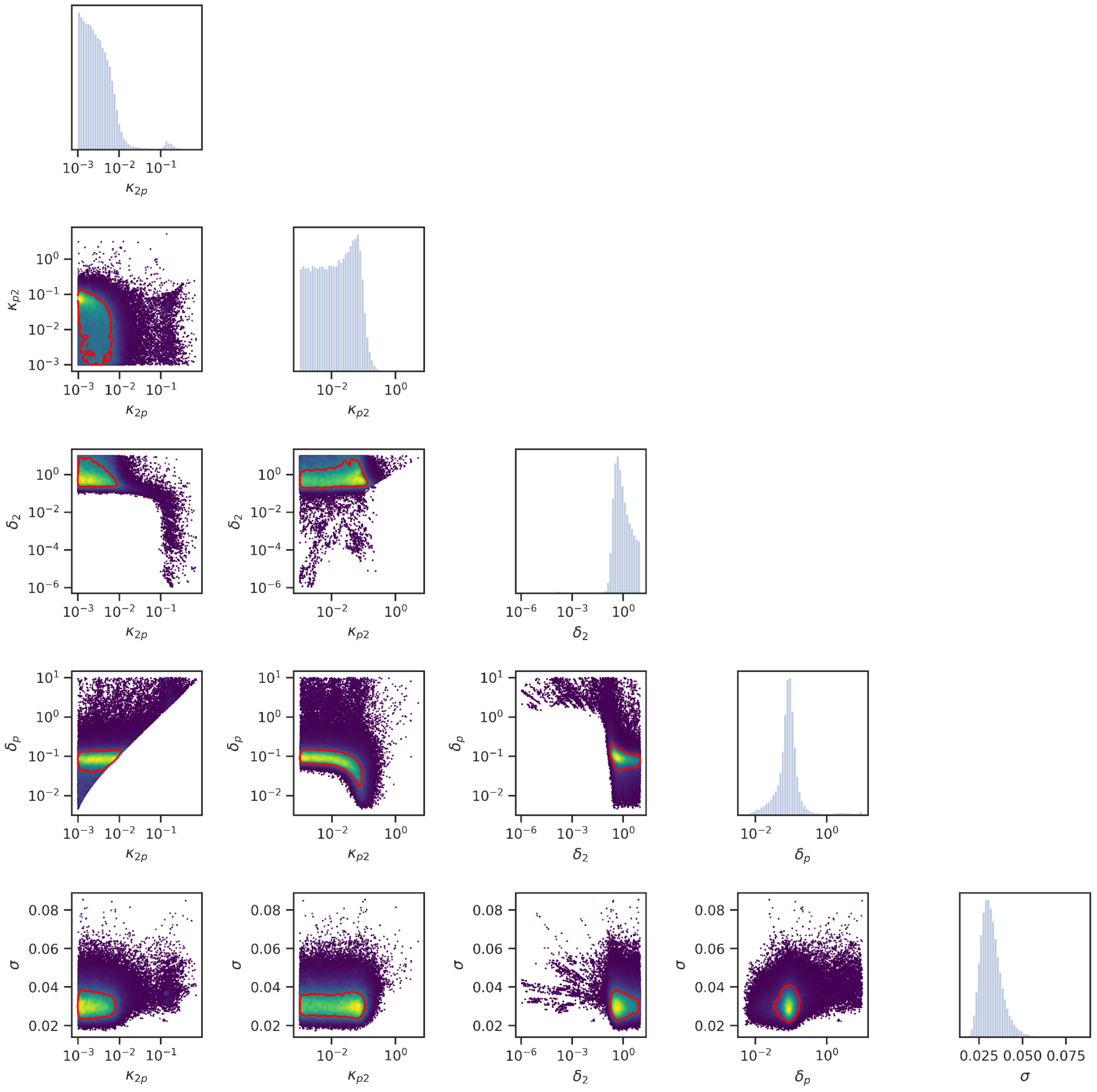


**Figure D: Posterior distributions of hepatocyte turnover parameters for the POP2p scenario estimated by MCMC sampling.**

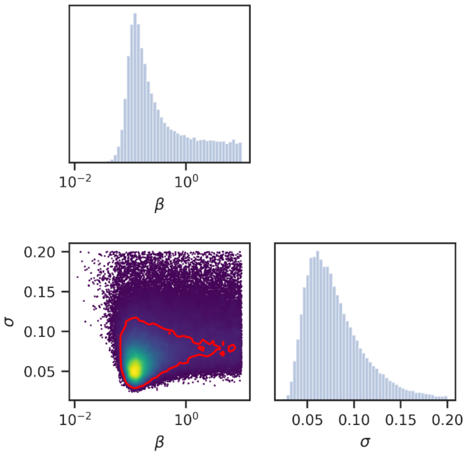


**Figure E: Posterior distributions of nonhepatocyte turnover parameters for the POP1 scenario estimated by MCMC sampling.**

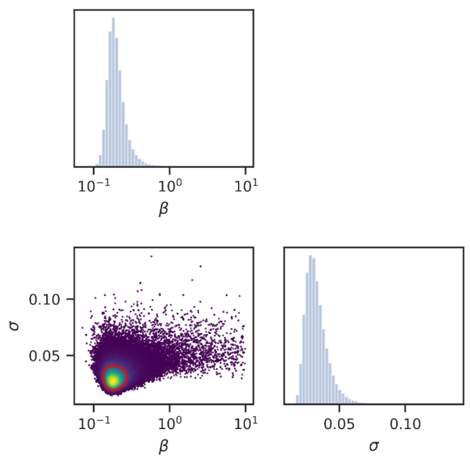


**Figure F: Posterior distributions of nonsorted liver cell turnover parameters for the POP1 scenario estimated by MCMC sampling.**
